# Allosterically coupled conformational dynamics in solution prepare the sterol transfer protein StarD4 to release its cargo upon interaction with target membranes

**DOI:** 10.1101/2023.03.24.534181

**Authors:** Hengyi Xie, Harel Weinstein

## Abstract

Complex mechanisms regulate the cellular distribution of cholesterol, a critical component of eukaryote membranes involved in regulation of membrane protein functions directly and through the physiochemical properties of membranes. StarD4, a member of the steroidogenic acute regulator-related lipid-transfer (StART) domain (StARD)-containing protein family, is a highly efficient sterol-specific transfer protein involved in cholesterol homeostasis. Its mechanism of cargo loading and release remains unknown despite recent insights into the key role of phosphatidylinositol phosphates in modulating its interactions with target membranes. We have used large-scale atomistic Molecular dynamics (MD) simulations to study how the dynamics of cholesterol bound to the StarD4 protein can affect interaction with target membranes, and cargo delivery. We identify the two major cholesterol (CHL) binding modes in the hydrophobic pocket of StarD4, one near S136&S147 (the Ser-mode), and another closer to the putative release gate located near W171, R92&Y117 (the Trp-mode). We show that conformational changes of StarD4 associated directly with the transition between these binding modes facilitate the opening of the gate. To understand the dynamics of this connection we apply a machine-learning algorithm for the detection of rare events in MD trajectories (RED), which reveals the structural motifs involved in the opening of a front gate and a back corridor in the StarD4 structure occurring together with the spontaneous transition of CHL from the Ser-mode of binding to the Trp-mode. Further analysis of MD trajectory data with the information-theory based NbIT method reveals the allosteric network connecting the CHL binding site to the functionally important structural components of the gate and corridor. Mutations of residues in the allosteric network are shown to affect the performance of the allosteric connection. These findings outline an allosteric mechanism which prepares the CHL-bound StarD4 to release and deliver the cargo when it is bound to the target membrane.

## INTRODUCTION

Cholesterol, a critical component of mammalian cell membranes, is heterogeneously distributed among cellular organelles. The involvement of Cholesterol (CHL) in the regulation of membrane protein function occurs both through direct protein-CHL interactions and through the effect of CHL on the physiochemical properties of the host membranes. Proper subcellular distribution of CHL is essential for membrane trafficking and cell signaling (1–4). The plasma membrane (PM) and the endocytic recycling compartment (ERC) are major pools of the total cellular cholesterol (1,5–8). On the other hand, the endoplasmic reticulum (ER), where the cholesterol concentration is sensed and regulated, and the cellular sterol homoeostasis is maintained (1,8,9), contains only a low amount of cellular cholesterol (0.1-2%).

Subcellular distribution of CHL occurs through both vesicular and non-vesicular transport mechanisms, the latter accounting for 70% of the cholesterol transport (5,8,10). Rapid non-vesicular transport requires lipid transfer proteins that provide the hydrophobic environment needed to accommodate lipid transferring across the aqueous phase (11,12). One major family of sterol transfer proteins is the (START) domain family of the steroidogenic acute regulatory protein (StAR)-related lipid-transfer. A START domain (STARD) is composed of ∼210 amino acids that fold into an α/β helix-grip structure to create an internal hydrophobic cavity for lipid binding and recognition as illustrated in Fig. 1A (13,14). The mammalian STARD protein family comprises 15 proteins, subdivided into six subfamilies based on domain architecture and ligand specificity (6,12,15,16). The characteristic of the STARD4 subfamily is a single soluble sterol-binding START domain. The StarD4 protein is well-adapted to bind and transport sterol (17,18) and is widely expressed in multiple tissues (19–21). StarD4 knockdown was shown to result in a decrease of (i) ER cholesterol concentration, (ii) acyl-CoA:cholesterol acyl-transferase 1 (ACAT1) activity, and (iii) cellular cholesterol ester concentration (7). The non-vesicular sterol transport kinetics between PM, ERC and ER was reduced by 33% (22). Conversely, StarD4 overexpression resulted in facilitated intracellular cholesteryl ester accumulation in a ACAT1 dependent manner (6,23).

**Figure 1:**
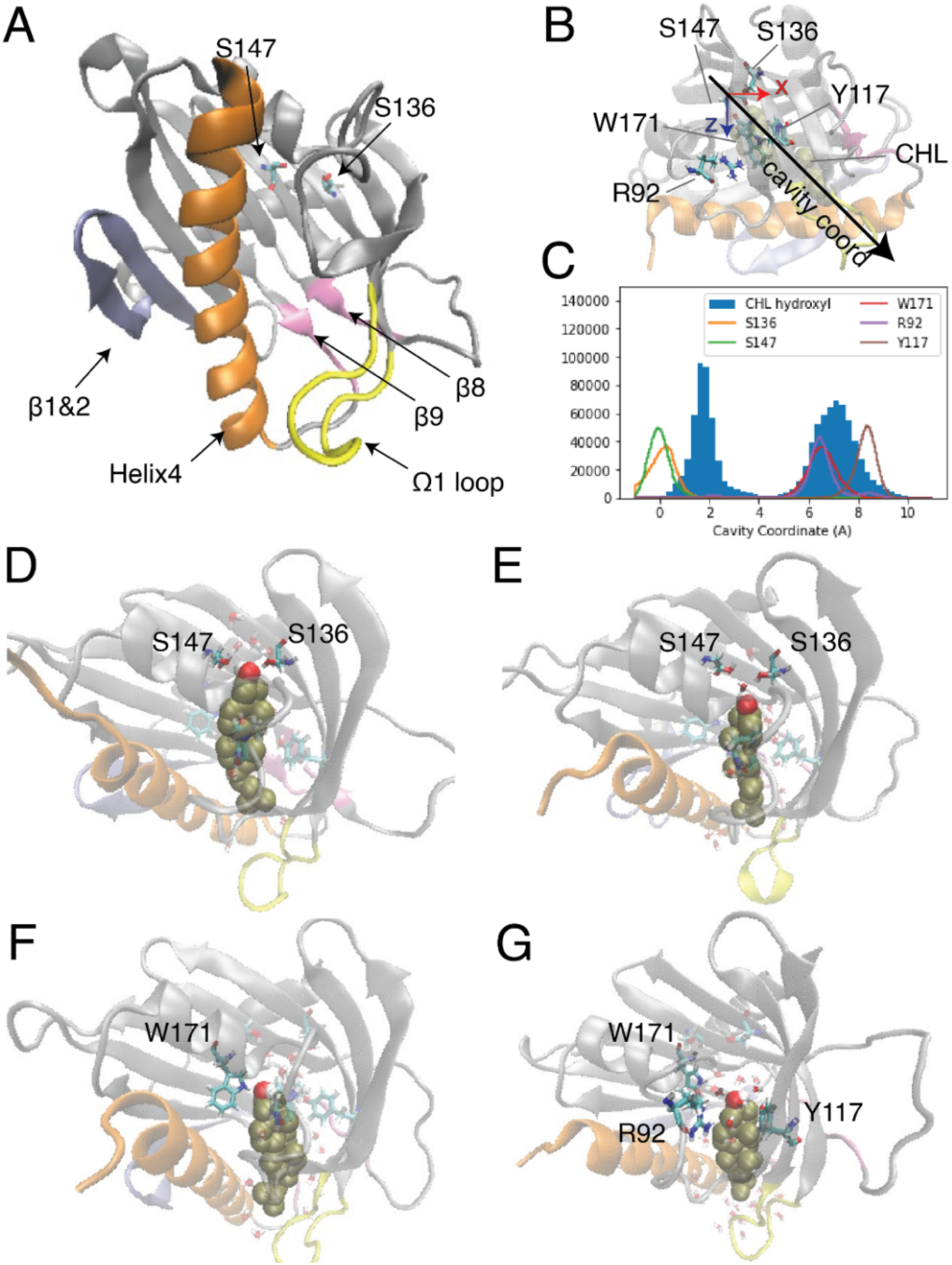
Two modes of CHL binding in the hydrophobic pocket of StarD4. The binding of cholesterol was modeled computationally in the crystal structure of StarD4 (PDB ID: 1jss) **(A)** The α/β helix-grip structure of the STARD domain in the crystal structure of StarD4. StarD4 is rendered in gray, with the C-terminal Helix (Helix4) in orange, the loop between β5 and β6 (Ω1 loop) in yellow, the β1&2 sheets in blue, and the β8 loop and β9 loop in pink. Previously reported potential cholesterol binding sites, residues S136 and S147, are rendered in “licorice” which draws the atoms as spheres and the bonds as cylinders, with oxygen colored in red, nitrogen in blue, carbons in cyan and hydrogens in white. **(B,C)** The “cavity coordinate” of CHL (rendered in VDW in tan color) and StarD4 residues reveal two modes of interaction. **(B)** Definition of the “cavity coordinate”: With the CHL-StarD4 complex rotated first to align the H4 along the x coordinate, and position the Center of Mass (CoM) of the protein right above the CoM of H4, the “cavity coordinate” is defined by taking the center of mass of S136&S147 as the origin and measuring the distance along the direction of the angle bisector of x- and z-axes. **(C)** Frequency histogram of the cavity coordinate of the hydroxyl oxygen in CHL (filled), and of the sidechain oxygen or nitrogen atoms in S136, S147, W171, R92 and Y117 (hollow). **(D-G)** Binding modes of CHL in the CHL-StarD4 complex. StarD4 is shown in the same representation as in (A). CHL is rendered in VDW with the hydroxyl group in red and white, and other atoms in tan color. Water molecules within 6Å from CHL are shown in transparent licorice rendering, and waters directly involved in hydrogen bonds with CHL and binding-sites are shown in opaque licorice. **(D)** the “Ser-binding mode”; **(E)** the “water-bridged Ser-binding mode”; **(F)** the “Trp-binding mode”; **(G)** “water-bridged Trp-binding mode”.

Despite extensive studies of STARD protein function, no structure of a START domain complexed with a sterol has been resolved. Thus, the ligand binding modes of the START domain remain undetermined, and the molecular mechanism that underlies the lipid specificity and determines the lipid uptake and release pathways, is still unclear. Studies of sterol binding in START domains that employed primarily docking and short MD simulation have identified the cholesterol binding pocket and residues likely to be involved in ligand binding (Fig. 1A) (14,24–27). In these studies, sterol was reported to favor a binding mode where the CHL inserts into the depths of the hydrophobic cavity and the hydroxyl group forms interactions with polar side chains or backbone carbonyls of residues at the bottom of of the pocket. In Fig. 1A residue S136 and S147 are labeled in the binding pocket because they correspond to the predicted CHL binding sites in StarD3 (S362 and R351) and StarD5 (S132).

The recently determined structures of LAM proteins in the StARkin superfamily show that the sterol hydroxyl group makes hydrogen bonds with residues located in the upper two-thirds of the hydrophobic tunnel, and the sterol hydrocarbon tail is exposed to the solvent through a partially open lid (28–30). Structural comparison between yeast LAM and human StarD4 also suggested that conformational changes at Helix4 and Ω1 loop (the loop between β5 and β6) (Fig. 1A) are required for the apo-StarD4 crystal structure to accommodate a cholesterol ligand at the same binding site as LAM protein (27).

In the present study we have used extensive (∼0.4 milliseconds) MD simulation trajectories to explore the binding modes of cholesterol in the pocket of StarD4 in solution in order to (1) assess the structural relationships between the *apo* and *holo* states of the protein, (2) identify the dynamic rearrangements required to accommodate the binding of CHL, and (3) the relation of these dynamic changes to the formation of the protein state required for its functionally productive membrane interactions. The long trajectories revealed a rich dynamic landscape of the protein structure in which the bound CHL adopts positions and configurations suggesting preparations for its CHL trafficking functions. Such function-related conformational changes of the StarD4 protein and its complex with CHL were revealed from the analysis of the MD trajectories with a machine learning-based Rare Event Detection (RED) protocol (31). To reveal the mechanisms underlying the function-related conformational changes we explored the allosteric pathways connecting the CHL repositioning in the binding site, with the identified conformational changes by applying the N-body Information Theory (NbIT) analysis (32,33) to the MD trajectories. We show here that the allosteric mechanism resulting from this analysis involves the coupling between the cholesterol translocation dynamics in the binding site and configurational changes of specific regions of the structure that are involved in the interaction of Stard4 with the membrane (34). The detailed structure-based information about the conformational changes and the allosteric channel enabled the development of specific testable hypotheses for mutations that we used here to probe the molecular mechanisms of function of a CHL-loaded StarD4 diffusing in the aqueous medium of the cytosol, and mutations that affect the molecular rearrangements that prepare the pathway for sterol release when the protein is embedding in a target membrane as we have shown (34).

## RESULTS

### MD SIMULATIONS REVEAL TWO MODES OF CHOLESTEROL BINDING IN StarD4

The binding modes of CHL in the pocket identified previously (14,16) were explored with extensive atomistic MD simulations comprised of 12 statistically independent replicates of 33.6 μs each for a total of 403 μs, starting from the crystal structure of StarD4 (PDBid: 1JSS) with a CHL molecule docked in the binding site using the Schrodinger Induced Fit Docking protocol (35,36) (see Methods for details). The simulations produced two different modes of CHL binding in the hydrophobic pocket (Fig. 1B), one with the hydroxyl group located near Ser136/Ser147 (termed “Ser-binding mode”), the other with the CHL OH near Trp171 (“Trp-binding mode”). In the “Ser-binding mode”, the cholesterol hydroxyl group forms hydrogen bonds with the Ser136 and Ser147 sidechains either directly, or through a water bridge (Fig. 1C,D). In the “Trp-binding mode”, the cholesterol hydroxyl group interacts either directly with Trp171, or resides between Trp171, Arg92 and Tyr117 with which it interacts through H-bonds mediated by a water molecule (Fig. 1E,F). Spontaneous transitions of cholesterol between binding modes are observed in more than half of the trajectories (Sup. Fig. 1).

### TRANSITIONS BETWEEN THE TWO MODES OF CHOLESTEROL BINDING ARE CONCURRENT WITH CONFORMATIONAL CHANGES OF THE StarD4 PROTEIN

To reveal the sequence of conformational events that accompany the transitions between the Ser and Trp binding modes, we applied the recently developed Rare Event Detection (RED) protocol (31) as described in Methods. The mechanistically important state-to-state transitions of StarD4 in response to the translocation of ligand cholesterol (see ref. 31 for a discussion of the relation between rare events and function) were extracted from the dynamic information in an ensemble of 6 trajectory stretches (4∼8 μs each, 34.4 μs total) from different trajectory replicas in which a stable cholesterol transition from one binding mode to another was observed. This ensemble of 6 trajectories served as the input training data for the RED protocol in identifying the rearrangement events <u>shared</u> among the different independent replicas.

The RED algorithm decomposed the trajectories into 5 components based on the evolution of the residue-residue contact map over time (see Methods). As described in ref (31), each component represents a state of StarD4 in which a particular conformation becomes dominant, at a particular time in the trajectory. This time-ordered series of events in which different components dominate the structural changes encoded in the trajectory, is shown in Fig. 2A,B. Several components identified by the RED algorithm are observed to be dominant concurrently with the repositioning of cholesterol in the binding site (Fig. 2A&C, B&D). This is remarkable, because the information provided to the RED algorithm consists only of the contact matrices for each frame in the trajectory, without direct information about CHL and its position. The correspondence between the binding modes and particular RED components is illustrated in the time frames by the correspondence of the times when component A (“comp A” in Fig. 2A,B) is dominant, and CHL is in the Ser-binding mode of CHL (Fig. 2C,D). Moreover, as the dominance of the A component wanes and is replaced by that of comp-s B and C, CHL is in the Trp-binding mode. Thus, in every independent trajectory the transition events from comp A to either comp B or comp C coincide with the translocation of cholesterol (Traj0&3 shown in Fig. 2 A-D, and Traj1,4,6,7 in Sup. Fig. 2A). This suggests a mechanism of coupled dynamics in the cholesterol-StarD4 complex, connecting the structural rearrangements of the protein frame with the transition of CHL in the binding pocket. The states associated with the transition of CHL to the Trp-binding mode are of particular interest because in this positions the CHL is near a putative “exit gate” from the binding pocket.

**Figure 2:**
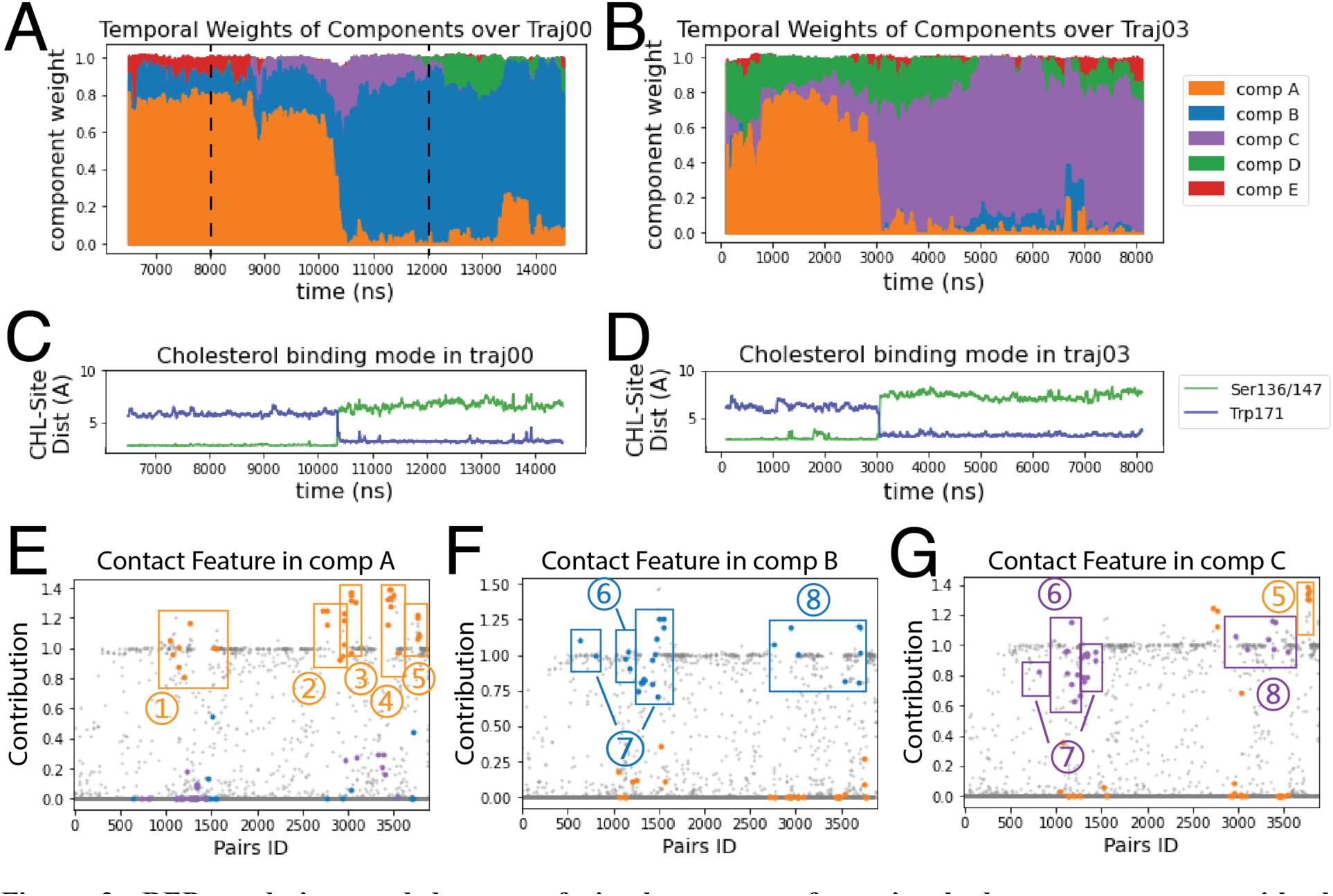
RED analysis revealed a set of simultaneous conformational changes concurrent with the translocation of CHL. **(A,B)** The time evolution of the RED-detected normalized temporal weight of structural components over the simulation time of trajectory 0 **(A)** and 3 **(B)**. Color code: comp A (orange), comp B (blue), comp C (purple), comp D (green), comp E (red). At each time point, the color column represents the stacked values of the weights for all the components beneath. For example, at time = 8000 ns in traj00 (left dashed line **in A**) the weight of comp A is 0.77, comp B is 0.13, comp E is 0.1, and comp C&D are 0; at time = 12000 ns (right dashed line), the weight of comp A is 0.05, comp B is 0.90, comp C is 0.03, comp D is 0.01 and comp E is 0. **(C,D)** The time evolution of the cholesterol binding mode, represented by the change in the distances between CHL and the Ser binding site (in green) and to the Trp site (blue) for: **(C)** in trajectory 0; **(D)** in trajectory 3. **(E-G)** Spatial arrays of components A, B, and C show the contact features characterized by the structure-differentiating contact pairs (SDCPs). Along the X axis there are 3880 data points, each representing one residue pair. The Y coordinate of each data point shows the contribution of the residue pair to the contact feature of the component (see Methods). The SDCPs (summarized in Table 1) are grouped by secondary structure location and highlighted in colors. Each figure (**E-G**) shows a conformational state for which the determinant conformational features are encoded by the contact patterns of structure-differentiating contact pairs (SDCPs). For example, in comp B (in **F**) residue pair groups 1-5 lost contact, while groups 6-8 have formed contacts. Together, these groups contain the SDCPs that define the structural feature in the time segment when comp B is prevalent in determining the conformational state.

We propose that the specific structural rearrangements of the protein frame constitute a preparatory step for a subsequent functional docking of the loaded StarD4 to the membrane for the release of its cargo. A key indication for the likelihood of this mechanism is the consistency in the different simulation replicas of the conformational changes coupled to the transitions between CHL binding modes.

#### RED analysis reveals the specific conformational changes that occur simultaneously

The nature of the conformational changes of the protein that are coupled to the CHL translocation toward the vicinity of the “gate” is identified from a comparison of the spatial arrays in the RED analysis.

For the analysis of the simultaneous conformational changes that constitute the rare event, we identified structure-differentiating contact pairs (SDCPs) among the large number of residue contacts included in a component (see protocol in Methods). A contact pair is deemed an SDCP if it satisfies two simultaneous criteria: the pair makes a substantial contribution to the contact feature in one component and a minimal contribution in another component. Both the SDCPs making significant contributions, and those making minimal contributions, are essential elements of a particular component feature, as they represent the formation and the elimination of residue pair contacts, respectively. They are essential in interpreting the RED analysis, because they collectively determine the conformational change characteristic to the rare dynamic event.

The SDCPs in Fig. 2E-G are grouped (from 1 to 8) by their location in a particular secondary structure motif. Specifically, group 1 includes contacts between β1 β2 & the C-terminal Helix (H4); group 2 includes contacts between Ω1 & H4 (Fig. 3A), group 3 includes contacts between Ω1 & β9 (Fig. 3A), group 4 has the contacts between β9 & β8 (Fig. 3C), and group 5 includes the interactions in the 1^st^ turn of H4 (Sup Fig. 4A).

**Figure 3:**
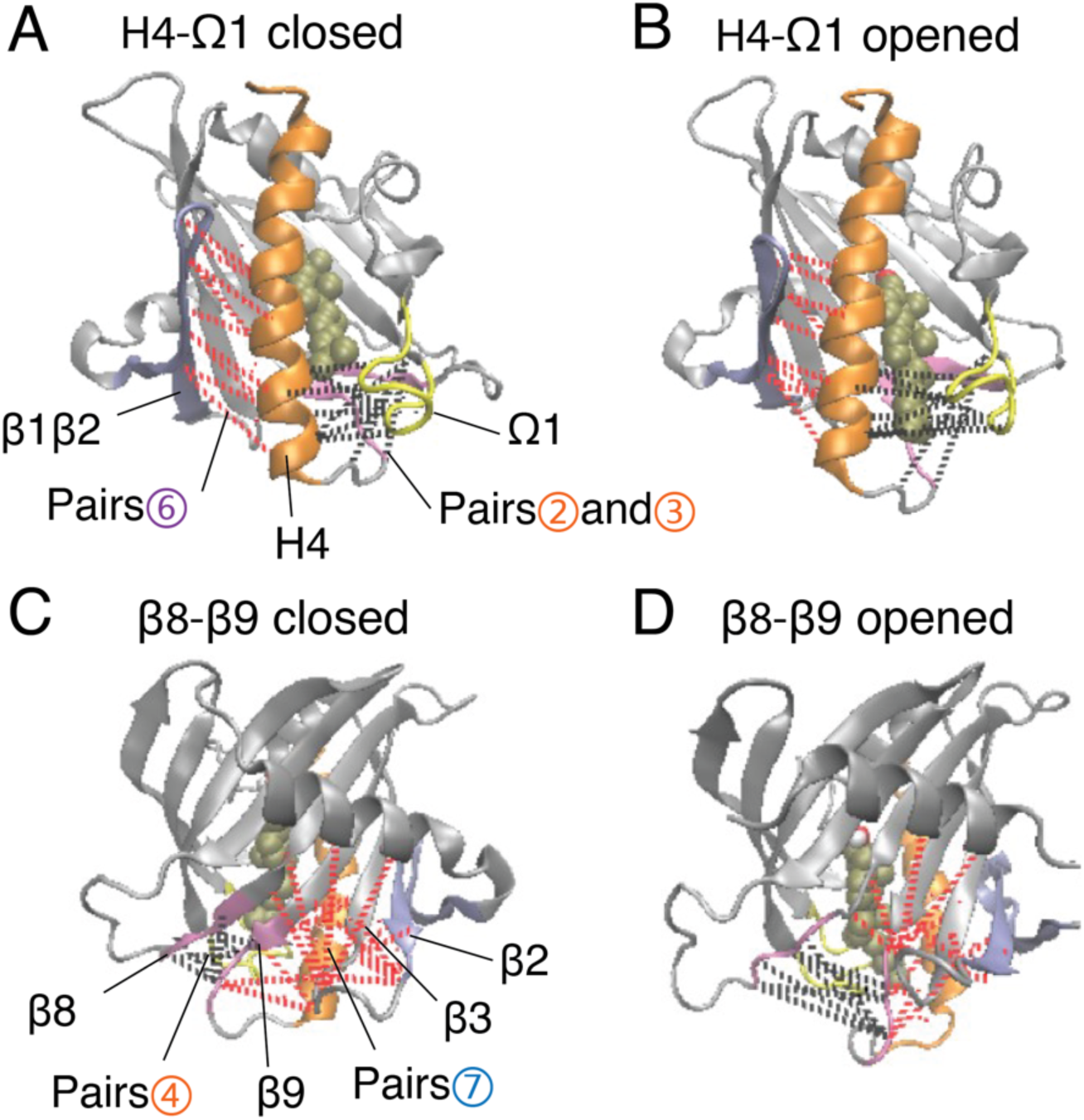
The translocation of CHL is concurrent with the opening of the gate and corridor to the hydrophobic pocket. **(A,B)** The elements of motif movements at the front gate between H4-Ω1 are compared in representative conformation in the closed state **(A),** and in the opened state **(B)**. The SDCPs containing breaking contacts (groups 2&3) are labeled by dashed lines in black, and those forming contact pairs (group 6) are labeled by red dashed lines. The representation and color scheme are the same as Fig. 1A. **(C,D)** The motif movements at the back corridor between β8-β9 are shown comparatively as the representative conformation in the closed state **(C)** and the opened state **(D)**. The contact breaking SDCPs (group 4) are labeled by dashed lines in black, and the contact forming SDCPs (group B7) are labeled by the red dashed lines.

The complete list of SDCPs is given in in Table1, below, which summarizes the structural characteristics of the system at different stages in the trajectory. The structural analysis shows that SDCPs in groups 1 to 5 are in contact in comp A in which the CHL is in the Ser-bound mode (orange, Fig. 2E), but most of them break contact during the transition from comp A to comp B or C, and thus have low feature values in comp B and comp C where CHL is in the Trp-binding mode (orange, Fig. 2F,G).

Another set of SDCPs, 6 to 8, have low values in comp A (blue and purple in Fig. 2E) but high values in comp B and C (blue in Fig. 2F and purple in Fig. 2G). They are found to represent contacts formed only <u>after</u> the repositioning of the CHL. More generally, comp B and comp C share a highly similar (but not identical) set of structural specifics represented in the salient SDCPs in groups 6 to 8. The highly similar pattern of motif interactions represented by these shared SDCPs include the following: in group 6, interactions between β1 β2 & H4 (Fig. 3B), in group 7, interactions between β2 & β3 & β9 (Fig. 3D), in group 8 interactions of β8 or β9 with Ω1 or H4. Thus, the summary in Table 1 shows that groups 6-to-8 in comp B (B6-B8) and groups C6-C8 in comp C share most of the key secondary structural elements and their conformations.

**Table 1.** Summarization of the structure-differentiating contact pairs constituting secondary structure motif interactions captured by comp A, B and C

| **Pair Group** | **Motif Interaction** | **Highly contact in** | **Residue Pairs** |
| --- | --- | --- | --- |
| 1 | Interactions between $\beta 1$, $\beta 2$ & H4 | comp A | R46-P60, V47-K50, K49-S208, K49-A211, D53-G73, V56-A211, Y67-V202, Y69-A207 |
| 2 | Contacts between $\Omega 1$ & H4 | comp A | Q122-A205, L123-A201, L123-T204, I126-A201, I127-A201, I127-V202, I127-A205 |
| 3 | Contacts between $\Omega 1$ & $\beta 9$ | comp A | I127-L193, R130-L193, R130-R194, R130-G195, R130-I197, F132-L193 |
| 4 | Contacts between $\beta 9$ & $\beta 8$ | comp A | V162-L193, V162-R194, R163-R194, G164-D192, G164-L193, G164-R194, Y165-R194, C169-W171 |
| 5 | Spiral contacts at the 1st turn of H4 | comp A & C | P198-A201, P198-V202, Q199-D203, S200-D203, S200-T204 |
| B6 | Shifted interactions between $\beta 1$ $\beta 2$ & H4 | comp B | K49-T204, V51-A207, V51-S208 |
| C6 | $\beta 9$ interactions with $\Omega 1$ or H4 | comp C | A48-D203, K49-D203, K49-T204, V51-A207, V51-S208, K52-A211, V54-L210, V56-M206, R58-Q199 |
| B7 | interactions between $\beta 2$, $\beta 3$ & $\beta 9$ | comp B | T33-F64, Y37-F64, R58-Y67, K59-Y67, P60-Y67, S61-G66, S61-Y67, S61-L68, S61-Q190, F64-H167, F64-P168, N65-D192, N65-R194, G66-R194, Y69-T191, Y69-L193 |
| C7 | Contacts between $\beta 9$ & $\beta 8$ | comp C | Y37-F64, R58-Y67, K59-Y67, P60-G66, P60-Y67, S61-G66, S61-Y67, S61-L68, S61-Q190, F64-H167, F64-P168 |
| B8 | $\beta 9$ interactions with $\Omega 1$ or H4 | comp B | L123-I197, I127-M196, R130-M196, P168-Q190, L193-G195, L193-M196, L193-M206, R194-M196 |
| C8 | $\beta 8$ interactions with $\Omega 1$ or H4 | comp C | S128-V162, V133-T157, W154-T157, R158-R163, P159-R163, E160-R163, P168-Q190 |

The major difference between comp B and comp C is the partial unfolding of the N-terminal head of H4 (group 5; residues 199-201) which loses some contacts in comp B but remains structured in comp C. Together, the structural features of components B or C compared to those of comp A reveal the specific conformational changes that are captured as “rare events”.

The main rare events identified by the transition from a conformation in which one component is dominant to a new conformation with a different dominant component, are the AèB and AèC transitions which are concurrent with the CHL translocation (Fig. 2A-D). The comparison of the structural features in comp A and in comp-s B&C shows that the conformational changes concurrent with the CHL translocation are the opening the access to the hydrophobic pocket. These conformational changes include (1) the opening of a front gate by the distancing of the H4 N-terminus from the Ω1-loop and strengthening interactions with β1 (see Fig 3A&B, and the detailed residue interaction data in Fig. 8A,B); (2) the opening of a back corridor between the β9 tail and the β7β8-loop (Fig 3C,D).

### THE PREDOMINANT CHANGES IN THE CONFORMATIONAL SPACE OF StarD4 EMERGE FROM CONCURRENT DYNAMICS OF CHL BINDING MODE SWITCHING AND GATE OPENING

To facilitate analysis of the metastable states and transition pathways encoded in the MD trajectory data, we built the conformational landscape of the cholesterol-bound StarD4 by applying the dimensionality reduction approach tICA: time-structure based independent component analysis. To describe the dynamics of cholesterol binding modes and of the protein conformational changes (see Methods and Sup. Table 1) we chose a set of 7 collective variables (CVs) to define the tICA space. The analysis showed that the first 3 tICA vectors describe 81% of the total dynamics of the system in the 403 μs production run trajectories (Fig. 4A,B). The projections on the space spanned by these vectors are shown in Fig. 4C.

**Figure 4:**
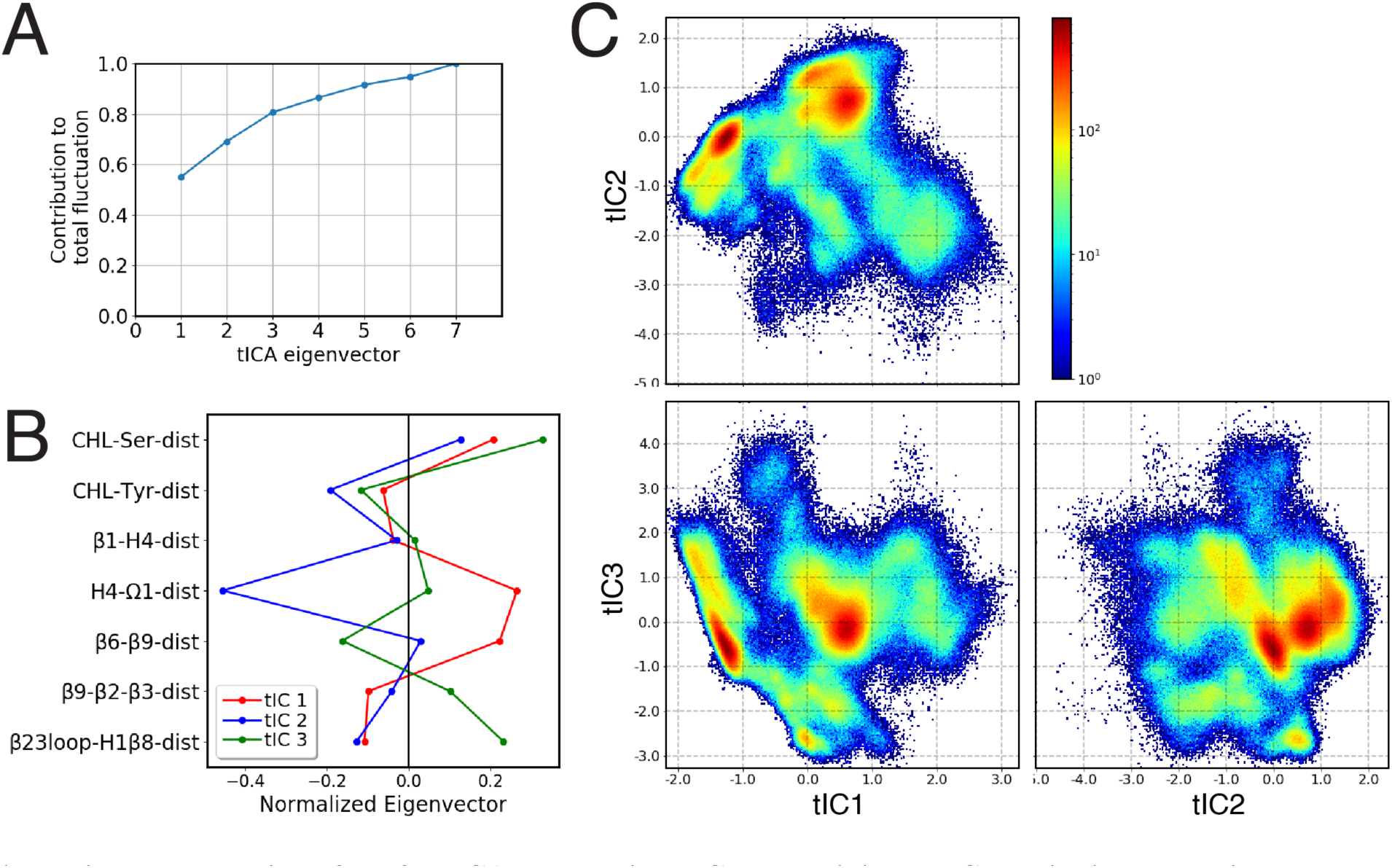
Representation of the 3D tICA space built on CVs describing the CHL binding modes in the pocket, and the protein conformational changes detected with the RED analysis. **(A)** Contribution of tICA eigenvectors to the total variations of the system contributed, showing that 81% of the information is captured by the first three tICA eigenvectors. **(B)** Contributions of individual CVs (which are identified on the Y axis, and defined in Sup. Table 1) to the first three tICA eigenvectors: tIC 1 (red), tIC 2 (blue), tIC 3 (green). **(C)** 2D projections of the 3D space defined by the 3 tICA eigenvectors, shown as population density maps: tIC2-tIC1 (top left), tIC3-tIC1 (bottom left), tIC3-tIC2 (bottom right). The population densities are colored according to the code shown on the right of the first panel.

The resulting 3D tICA space was discretized into 200 microstates using k-means clustering, and the transitions among these states were analyzed with the Markov State Model (MSM) approach (see Methods and Sup. Fig. 5). By segregating the 200 microstates according to kinetic similarities we identified 9 macrostates. The 5 macrostates with high population are shown in Fig. 5A, and representative conformations obtained from structural analysis are shown in Figs. 5B-D.

**Figure 5:**
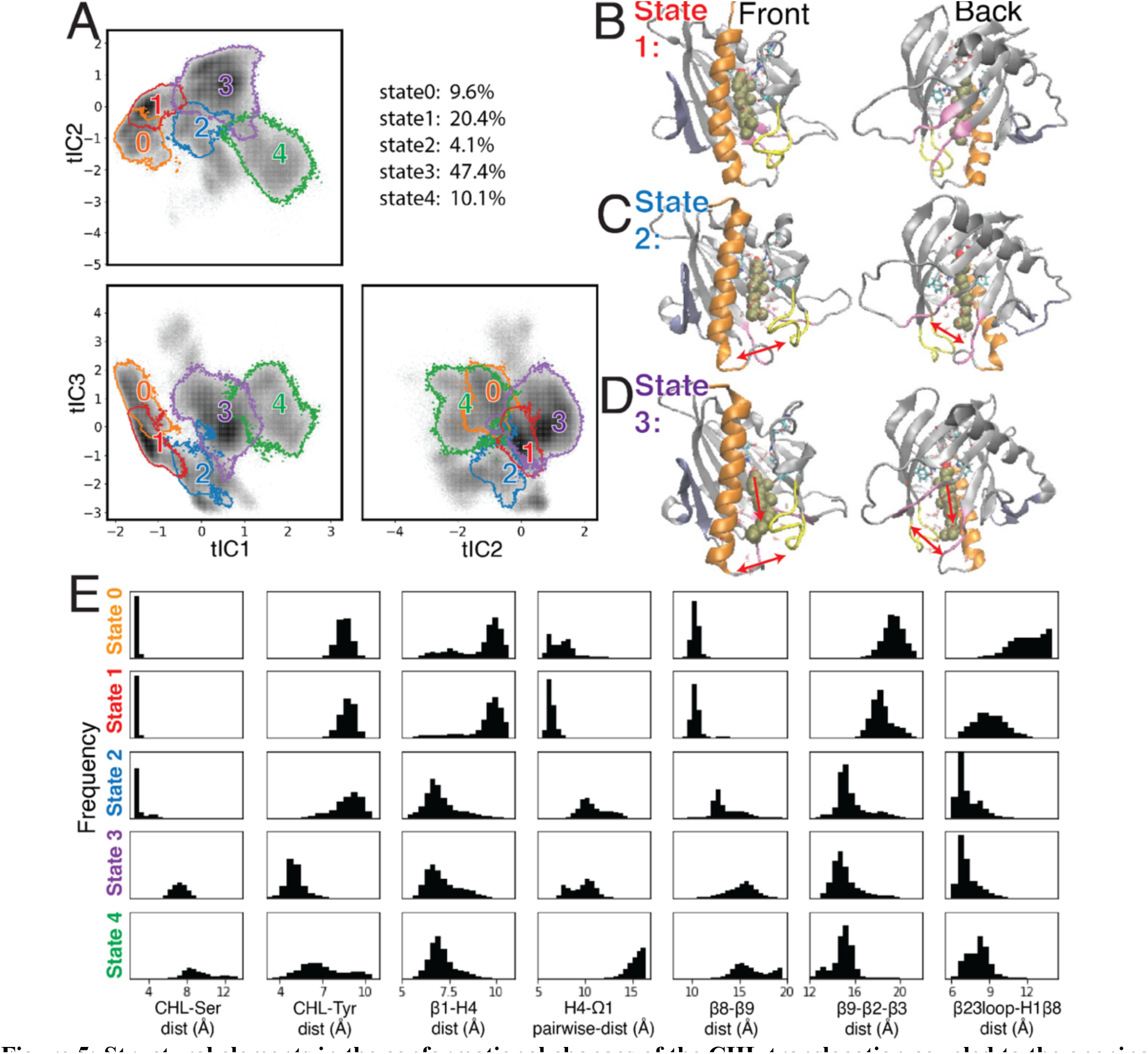
Structural elements in the conformational changes of the CHL translocation coupled to the opening of the H4-Ω1 gate and β8-β9 corridor. **(A)** Macrostates 0-4 are outlined in the 2D projections of the 3D conformational space: the tIC2-tIC1 plane (upper left), tIC3-tIC1 plane (lower left), tIC3-tIC2 plane (lower right). The gray shade represents the population density in the conformational space as shown in Fig. 4C. The populations (% of total microstates sampled) are listed at the upper right for each microstate. **(B-D)** Structural models representing macrostates 1 to 3 viewed from front (left column) and back (right column). The StarD4-CHL complex is rendered in the same way as in Fig. 1. Red arrows in the 2^nd^ and 3^rd^ rows **(C and D)** indicate the conformational changes of the H4-Ω1 gate and the β8-β9 corridor, as well as the movement of CHL. **(E)** Structural characteristics of macrostates 0 to 4 shown represented by probability density histograms of the characteristic CV values that determine the tICA space, as defined in Sup. Table 1.

The characterization of macrostates 0-4 by the relevant CVs (Fig. 5E), reflects the concurrent dynamics of gate, corridor, and cholesterol we detected in the stable transition events. The data in Fig. 5E show that these relations prevail throughout the conformational space.

Macrostate 1 (Fig. 5B) represents conformational states where cholesterol is bound in the Ser-site mode (with low “CHL-Ser-dist” and high “CHL-Tyr-dist”, seen in the1^st^ and 2^nd^ panel of Fig. 5E), in crystal-like conformations of StarD4 with a closed H4-Ω1 gate (high “β1-H4-dist”, low “H4-Ω1-dist” in 3^rd^ and 4^th^ panel of Fig. 5E) in which the β8-β9 corridor is closed (low “β6-β9-dist” and high “β9-β2-β3-dist”&“β23loop-H1β8-dist”, 5^th^ to 7^th^ panel in Fig. 5E). Macrostate 0 also represents conformational states with Ser-binding CHL in crystal-like StarD4 with gates closed, but with a conformation of the β23loop that is extended away (Sup. Fig. 6A) (high “β23loop-H1β8-dist”, 7^th^ panel, Fig. 5E) which may hinder the opening of β8-β9 corridor. These Ser-binding/gates-closed conformations in Macrostates 1 and 0 contribute 30% of the populations in the conformation space.

The most populated macrostate is Macrostate 3, with 47% of the population. It contains conformational states in which CHL is in the Trp-binding mode with both the H4-Ω1 gate and β8-β9 corridor open, and with β9 leaning towards β2&β3 which rearrange into narrower conformations (Fig. 5D,E). All these observed rearrangements are consistent with the conformational changes detected by the RED analysis of the corresponding RED event. In Macrostate 4 the cholesterol is in a Trp-binding binding mode or even further down towards the H4-Ω1 gate that opens wider (Fig. 5E, Sup. Fig. 6B). As indicated by the analysis, Macrostates 0,1,3,4 constitute 88% of the conformational space containing the concurrent dynamic rearrangements of the cholesterol binding mode, the opening H4-Ω1 gate, and the β8-β9 corridor. The other structural motif movements that were most important in the transition event detected with RED are also evidenced by comparisons of CVs related to the transition between the Ser- and Trp-binding modes of CHL (detected by the transition for Component A to B/C in RED). These are quantified in Sup. Fig. 7 and include the unfolding in H4 as well as the straightening of H4, the narrowing of β2 and β3, and residue interactions between β1-H4 and disassociations between β8-β9. Thus, the conformational changes of the StarD4 protein that we found to occur simultaneously with the transitions between CHL binding modes, yield conformational states that are specific to the position of CHL in the binding pocket.

The time sequence of cholesterol translocation in the binding site is embedded in the transition pathways starting from Macrostate 1 (crystal-like StarD4 with Ser-binding cholesterol) and ending in Macrostate 3 (gates-opened StarD4 with Trp-binding cholesterol). The fluxes along the most probable pathways (contributing ∼95% of the transition flux) are summarized in Table 2. Each of the top two most probable paths contributes ∼30% of the total flux. One is the direct transition from Macrostate 1 to Macrostate 3 which encompasses the simultaneous translocation of CHL and conformational change of StarD4, and the other involves an intermediate, the less populated Macrostate 2 in which CHL is in Ser-binding modes but with both H4-Ω1 gate and β8-β9 corridor open (Fig. 5C, E). These results are consistent with the structural analysis in a report (27) showing that conformational changes at the H4-Ω1 gate are required for cholesterol to bind lower in the hydrophobic pocket, due to steric hindrance.

**Table 2.** Top transition pathways in the cholesterol translocation process from Ser-binding state (1) to Trp-binding state (3)

| Path | Norm Flux | Cumulative Flux |
| --- | --- | --- |
| 1 → 2 → 3 | 0.3269 | 0.3269 |
| 1 → 3 | 0.2984 | 0.6253 |
| 1 → 0 → 7 → 3 | 0.1151 | 0.7404 |
| 1 → 0 → 2 → 3 | 0.0826 | 0.8230 |
| 1 → 0 → 3 | 0.0464 | 0.8754 |
| 1 → 5 → 3 | 0.0414 | 0.9168 |
| 1 → 0 → 2 → 4 → 3 | 0.0258 | 0.9426 |
| 1 → 5 → 2 → 4 → 3 | 0.0140 | 0.9567 |

The less populated intermediate macrostates are summarized in Sup. Fig. 8. They indicate the flexibility of the StarD4-cholesterol complex and support the relation between CHL position and the gate and corridor dynamics. Thus, with CHL in the Ser-binding mode, the gates-closed crystal-like conformations represent 30% of the population, much more than the ones with either the H4-Ω1 gate or the β8-β9 corridor open (10%). Most likely this is due to the energy cost of exposing the StarD4 hydrophobic pocket and cholesterol’s hydrophobic tail to water. In contrast, when CHL is in the Trp-binding mode the H4-Ω1 gate is always open, and the conformations with both gate and corridor open amount to 58% compared to those with β8-β9 corridor closed, which are very unlikely (1.7%).

### THE ALLOSTERIC PATHWAY COUPLING CHL TRANSLOCATION IN THE HYDROPHOBIC POCKET TO THE OPENING OF ACCESS GATES

Having identified a set of concurrent conformational changes in several structural motifs that include the cholesterol binding site, we applied NbIT analysis to reveal the allosteric network that connects them. NbIT analysis calculates the normalized coordination information (NCI), which describes how much information of the *receiver* system can be gained from information about the *transmitter* system (see ref 32 and Method for details). Here we applied the NCI analysis to the trajectories from the StarD4-cholesterol complex simulations to quantify the information transmission between motifs that emerged from the RED analysis described above. The residue index in Method identifies the components of the structural motifs used in this analysis summarized in Table 3. The structural motifs serving as negative controls for which the NCI quantification is expected to reveal only background-level information sharing with the cholesterol binding site are defined in Sup. Table 2.

**Table 3.**
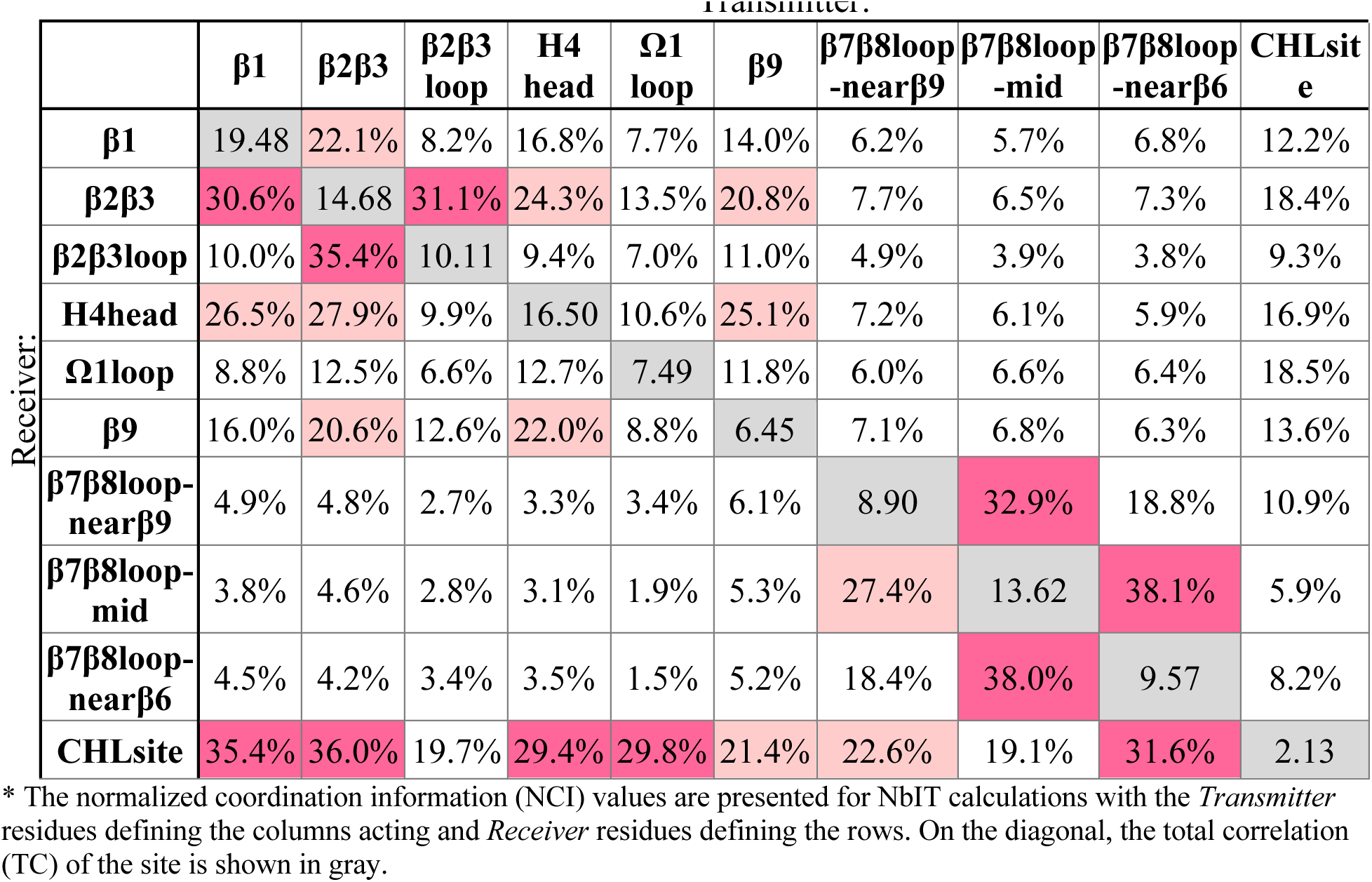
Normalized coordination information between sites in cholesterol-StarD4 system

|  |  | Transmitter: |  |  |  |  |  |  |  |  |  |
| --- | --- | --- | --- | --- | --- | --- | --- | --- | --- | --- | --- |
| Receiver: | | $\beta 1$ | $\beta 2\beta 3$ | $\beta 2\beta 3$ loop | H4 head | $\Omega 1$ loop | $\beta 9$ | $\beta 7\beta 8$ loop -near$\beta 9$ | $\beta 7\beta 8$ loop -mid | $\beta 7\beta 8$ loop -near$\beta 6$ | CHLsite e |
| | $\beta 1$ | 19.48 | 22.1% | 8.2% | 16.8% | 7.7% | 14.0% | 6.2% | 5.7% | 6.8% | 12.2% |
| | $\beta 2\beta 3$ | 30.6% | 14.68 | 31.1% | 24.3% | 13.5% | 20.8% | 7.7% | 6.5% | 7.3% | 18.4% |
| | $\beta 2\beta 3$ loop | 10.0% | 35.4% | 10.11 | 9.4% | 7.0% | 11.0% | 4.9% | 3.9% | 3.8% | 9.3% |
|  | H4head | 26.5% | 27.9% | 9.9% | 16.50 | 10.6% | 25.1% | 7.2% | 6.1% | 5.9% | 16.9% |
| | $\Omega 1$ loop | 8.8% | 12.5% | 6.6% | 12.7% | 7.49 | 11.8% | 6.0% | 6.6% | 6.4% | 18.5% |
| | $\beta 9$ | 16.0% | 20.6% | 12.6% | 22.0% | 8.8% | 6.45 | 7.1% | 6.8% | 6.3% | 13.6% |
| | $\beta 7\beta 8$ loop -near$\beta 9$ | 4.9% | 4.8% | 2.7% | 3.3% | 3.4% | 6.1% | 8.90 | 32.9% | 18.8% | 10.9% |
| | $\beta 7\beta 8$ loop -mid | 3.8% | 4.6% | 2.8% | 3.1% | 1.9% | 5.3% | 27.4% | 13.62 | 38.1% | 5.9% |
| | $\beta 7\beta 8$ loop -near$\beta 6$ | 4.5% | 4.2% | 3.4% | 3.5% | 1.5% | 5.2% | 18.4% | 38.0% | 9.57 | 8.2% |
|  | CHLsite | 35.4% | 36.0% | 19.7% | 29.4% | 29.8% | 21.4% | 22.6% | 19.1% | 31.6% | 2.13 |

\* The normalized coordination information (NCI) values are presented for NbIT calculations with the *Transmitter* residues defining the columns acting and *Receiver* residues defining the rows. On the diagonal, the total correlation (TC) of the site is shown in gray.

The results from the NCI calculation show that the β1, β2β3, H4 head, and Ω1 loop motifs are strongly coordinated with the cholesterol binding site (S136, S147, W171, R92, Y117), while weaker coordination is found with the β9 and the β7β8 loop regions. Notably, the H4 head and Ω1 loop constitute the H4-Ω1 gate, and the β9 and the β7β8 loop constitute the β8-β9 corridor of the hydrophobic pocket. The stronger coordination of the CHL binding site by the H4-Ω1 gate than β8-β9 corridor is in accordance with the structural analysis in the previous section showing that the conformational changes and interaction changes at the H4-Ω1 gate always accompany CHL translocation to the Trp-binding state (Fig. 2H,I)

Because β1 and β2β3 are also strong coordinators of the binding site despite the long distance separating these motifs, we calculated the *normalized mutual coordination information* (NMCI) to detect the coordination channel for information transmission from β1 to the CHL binding site (Fig. 6A). The calculated NMCI shows that β2β3, H4, and β9, all share the information with the binding site, which identifies them as components of the *coordination channel*. Thus, the coordination of cholesterol binding mode by β1 is shown to be mediated through the interaction of β1 with β2β3 and H4, which stabilizes the β9 and H4 in the gate opening conformation (Fig. 6B).

**Figure 6.**
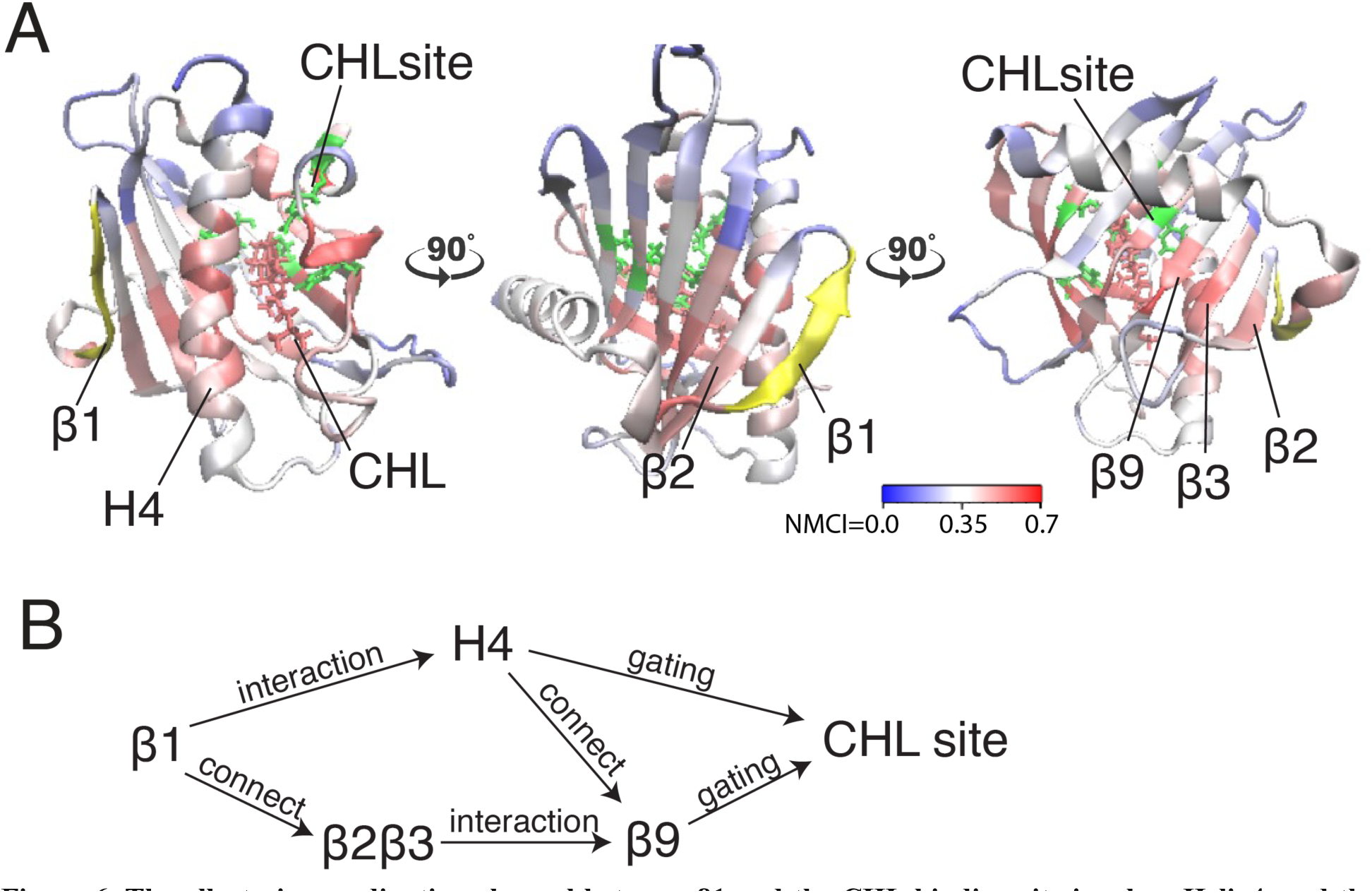
The allosteric coordination channel between β1 and the CHL binding site involves Helix4, and the β2β3 and β9 sheets. **(A)** StarD4 is rendered in carton, with residues colored according to the normalized mutual coordination information (NCMI) value calculated with the NbIT information theory-based analysis (32,33). The quantification scale is at the lower right. The *transmitter motif* (β1 residues 47-52) is colored in yellow, and the *receiver motif* (CHL binding site residues S136, S147, W171, R92 and Y117) is labeled in green cartoon on the secondary structure with side chains in green licorice. **(B)** An allosteric model of information transmission from β1 to the cholesterol binding site.

### THE PROBABILITY AND STABILITY OF CHL BINDING IN THE Trp-binding MODE ARE REDUCED IN THE W171A MUTANT StarD4

The release pathway of CHL in the yeast sterol transport protein *Osh*4 (37) was shown to comprise step-wise interactions of the CHL hydroxyl with a sequence of hydrophilic residues inside the hydrophobic pocket. In StarD4 we observed a similar sequence of CHL occupying binding sites along the pocket, composed of the S136&S147-site (Fig. 1D), a water-mediated S136&S147-site (Fig. 1E), the W171-site (Fig. 1F), and a water-mediated W171, R92 & Y117 site (Fig. 1G). Given the coordination between CHL translocation in the binding site and specific changes in the conformation of the StarD4 protein identified from the analysis of MD simulation trajectories, we probed the impact of impairing such a binding site by introducing a W171A mutation. We used a Free Energy Perturbation (FEP) protocol to introduce the mutation in representative conformations of the two most populated Microstates (Fig. 5): State 1 with CHL in a Ser-binding mode, and State 3 where CHL interacts with W171 and is also involved in the water bridged interactions with R92 and Y117. From each binding mode we launched 24 replicate runs of 2,720ns/each, resulting in an ensemble of simulations totaling 131μs for the W171A-StarD4 mutant.

In simulations starting from the Ser-binding state, we find that the W171A mutation eliminated the translocation of CHL away from the Ser-sites, as the ligand is either in the Ser-binding mode or in the water-bridged Ser-binding mode. In the simulations started from the Trp-binding-like state where the CHL is interacting with R92 and Y117 (through a water-bridge), the W171A mutation destabilizes this binding mode and the CHL tends to translocate from the Trp-binding-like state back to the Ser-binding site. The backward transitions are observed in 8 out of the 24 replicas, with the Ser-binding conformation contributing 21% of the population in the ensemble started from the Trp-binding-like mode (Sup. Fig. 9A). In comparison, stable backward transitions were observed only 2 times in the 403 μs simulations of the WT StarD4 (Sup. Fig. 1). Thus, W171 turns out to be an important intermediate site that enables accessibility and stability for CHL binding to the downstream binding sites around W171, R92 and Y117.

Notably, structural analysis suggests that the allosteric connection persists in the W171A StarD4-CHL complex and the conformation of StarD4 is compatible with the CHL in the Ser-binding mode. Thus, when CHL transitions back from the R92 Y117 water network to the Ser-binding site, the initially open Ω1-H4 gate closed, and the unraveled N-terminal of H4 folded back (Fig. 7 and Sup. Fig. 9B-D). These conformational changes are consistent with the allosterically connected changes in WT StarD4 where the Ω1-H4 gate opening and the unfolding in H4 are found to be dominant features in the RED analysis only in the Trp-binding mode (Fig. 5E and Sup. Fig. 7). Results from the NbIT analysis of the mutant confirm that the allosteric effect remains strong between the CHL and binding site to the peripheral transmitter motifs around H4 (Sup. Table. 4).

**Figure 7.**
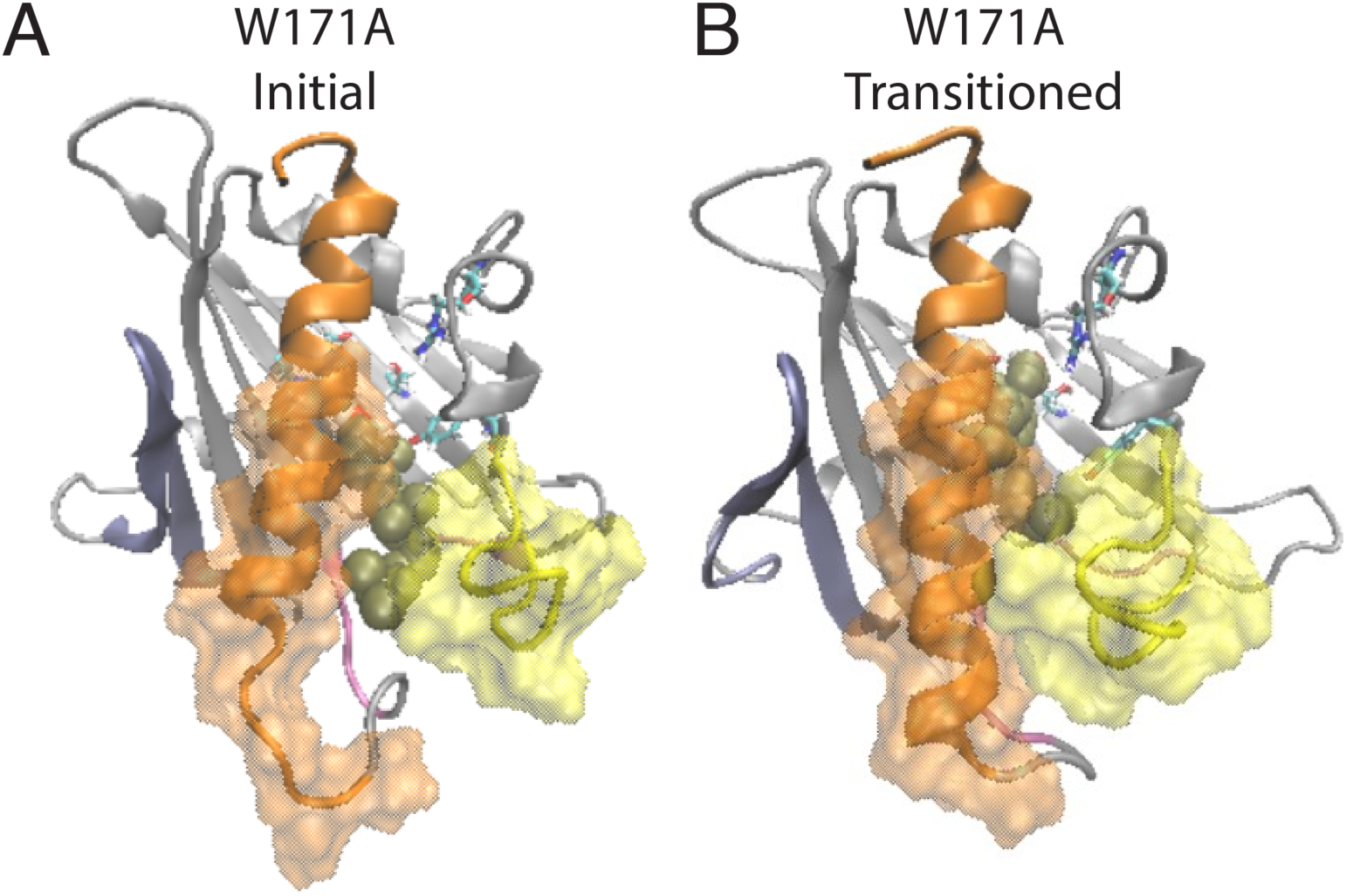
β1 and H4 conformational changes are allosterically coupled with the translocation of CHL binding in the StarD4 W171A mutant. **(A)** Starting position of the CHL in the simulation of the W171A StarD4 structural model. CHL resides in the water-bridged binding site with R92 and Y117. **(B)** Shows that the CHL has transitioned back up the hydrophobic binding site to the Ser-binding site in which the gate and corridor are closed. StarD4 is rendered in gray, with the H4 in orange, the Ω1 loop in yellow, the β1&2 sheets in blue, and the β8 loop and β9 loop in pink. Residues S136 and S147 are rendered in “licorice” which draws the atoms as spheres and the bonds as cylinders, with oxygen colored in red, nitrogen in blue, carbons in cyan and hydrogens in white. CHL is rendered in tan color, in the VDW representation. The surface of residues 198 to 212 of H4 is rendered in transparent orange, and the surface of residues 121 to 130 on Ω1 loop is rendered in transparent yellow.

### MUTATIONS OF K49 DISRUPT THE ALLOSTERIC NETWORK AND SLOW DOWN THE DYNAMICS OF CHL TRANSITIONS IN THE BINDING SITE

The mutation was chosen based on the results from analysis with the RED algorithm which revealed that the transitions of CHL in the binding site are coupled to a shift of the K49-S208 interaction to a K49-T204 configuration, and the establishment of a V51-A207 contact (Table 1). According to the allosteric model created with NbIT, β1 is an allosteric transmitter that coordinates the dynamics of bound CHL through its interaction with H4 and β2. The H4-β1 interaction involved in H4 repositioning is stabilized by both electrostatic and hydrophobic interactions. Thus, when the H4-β1gate is closed the K49 sidechain relocates near S208 and its hydrocarbon chain interacts with A207 (Fig. 8A). When the H4-β1gate opens, the K49 sidechain shifts closer to T204, enabling V51 to form a contact with A207 and remove this hydrophobic residue from an unfavorable aqueous environment (Fig. 8A,B). The NbIT coordination contribution analysis confirms that K49 is contributing the most to the allosteric interaction between β1 and the CHL binding site residues. Therefore, we mutated K49 expecting changes of the β1-H4 interaction mode that weaken the allosteric coordination.

Mutations K49A and K49W were introduced with the same FEP protocol as above into the StarD4-CHL complex in Microstates 1 (Ser-binding mode) and 3 (Trp-binding mode). Then, simulations were carried out for 24 replicas from each state for 2,320ns per replica of the K49A-StarD4 (111μs total), and 2,720ns per replica for K49W-StarD4 (131μs total). When started from the Ser-binding mode, the CHL transitioned to the Trp-binding site in 4 out of 24 trajectories for K49W-StarD4 (containing 7% of the population, Sup. Fig. 10A), and in only 1 out of 24 trajectories for K49A-StarD4 (containing 2% of the population, Sup. Fig. 10B). These results suggest that the dynamics of CHL are significantly slower in the mutants compared to the WT StarD4 in which 6 out of 12 trajectories sampled the transition within the first 2720ns. No backward transition from a Trp-binding state to a Ser-binding state was observed in the K49A and K49W mutant constructs. These results are consistent with decreased CHL dynamics in the K49W and in the K49A mutants.

To study the conformational changes and allosteric effect, adaptive sampling was carried out to expand the sampling along the reaction pathway of the K49A mutation for another 48 μs in 24 replicas (Sup. Fig. 10C). The impact of the mutation on the allosteric mechanism of conformational changes in StarD4 is evident in the change of contact probability between residues in β1 and H4 as shown in Table 4. Resulting structures compared in Fig. 8 show how the K49W and K49A mutations changed the mode of interaction between β1 and H4. In the WT StarD4, K49 establishes both hydrophobic and hydrophilic contact with A207, S208 and T204, and its translocation opens space for the formation of a V51-A207 contact pair (Fig. 8A,B). With the K49W mutation, the large W49 forms stable contacts with both S208 and T204, but hinders sterically the V51-A207 interaction (Fig. 8C and Table 4). The K49A mutation, on the other hand, resulted in a decreased contact rate with both S208 and T204, eliminated the hydrophilic interactions, and destabilized the V51-A207 contact (Fig. 8D and Table 4).

**Table 4.** Contact probability* between β1 and H4 in WT, K49W and K49A StarD4 WT:

| Contact probability | K49 – T204 | K49 – S208 | V51 – A207 |
| --- | --- | --- | --- |
| Ser-binding | 13.6% | 68.8% | 14.8% |
| Trp-binding | 83.3% | 13.1% | 75.4% |

**K49W:**
| Contact probability | W49 – T204 | W49 – S208 | V51 – A207 |
| --- | --- | --- | --- |
| Ser-binding | 51.0% | 77.1% | 1.3% |
| Trp-binding | 91.4% | 93.2% | 0.3% |

**K49A:**
| Contact probability | A49 – T204 | A49 – S208 | V51 – A207 |
| --- | --- | --- | --- |
| Ser-binding | 2.8% | 10.9% | 3.2% |
| Trp-binding | 76.5% | 17.7% | 39.6% |

\*Defined as the probability of the specified residue pairs having at least one atom in each at a distance <3.5Å. The Probability is expressed as the percent occurrences of such contacts.

**Figure 8.**
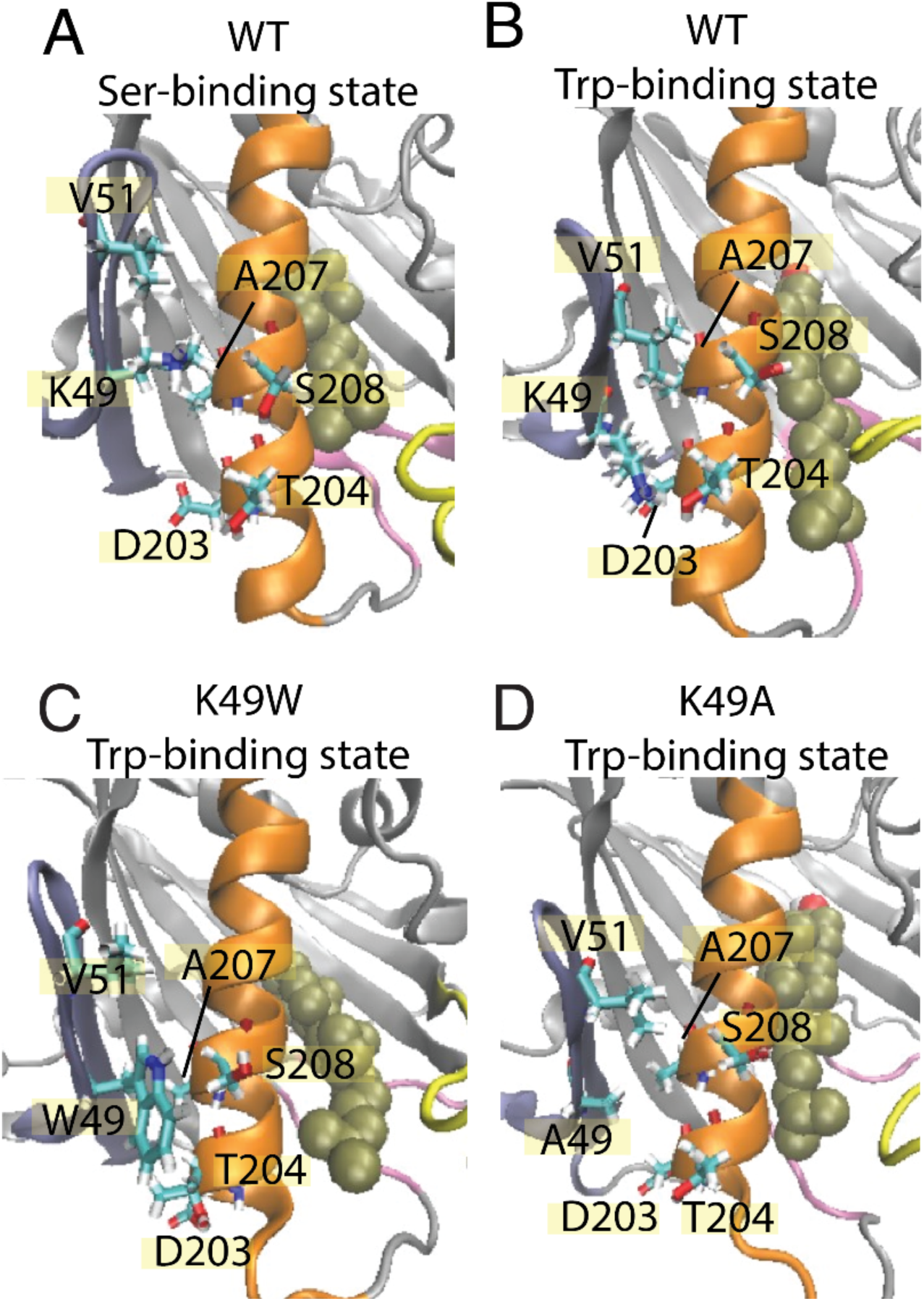
The β1-H4 interactions that are allosterically coupled with CHL translocation in WT are destabilized in K49 mutants. Comparison of detailed changes in H4 and β1 residue interaction modes in **(A)** WT StarD4 in the initial Ser-binding state; **(B)** WT StarD4 after transition to the Trp-binding state, **(C)** K49W StarD4 after transition to the Trp-binding state, **(D)** K49A StarD4 after transition to the Trp-binding state. The representation and color scheme are as in Fig. 1A. Residues K/W/A49, V51, D203 T204, A207 and S208 are rendered in “licorice”.

The interactions between β1 and H4 are weakened overall in both mutants but the interaction modes differ. The probability of contact is decreasing gradually from WT to K49W to K49A, and the electrostatic interaction is also gradually removed in the same order. Consistent with the weakening of interaction and the decrease of CHL dynamics, the NbIT results showed a decrease in coordination information (CI values) in the allosteric network between the β1, H4 and β9 motifs the K49W mutant (Sup. Table. 5), and a loss of coordination effect from β1 to the CHL site in the K49A mutant (Sup. Table. 6) – as expected from our results from the analysis of WT StarD4.

## DISCUSSION

The function of StarD4 in vectorial transport of sterols between preferred organelles is reported to be regulated by anionic lipids (34). To learn about the molecular mechanism that underlies the StarD4 sterol transport, and the role of phosphatidylinositol phosphate (PIP) anionic lipids in regulating its function, we explored the structural and dynamic properties of the protein and the binding modes of cholesterol (CHL). Our findings show that CHL can shuttle between two modes of stable binding in the hydrophobic pocket of StarD4, interacting as shown in Fig 1 either with Ser136 and Ser147 which are located at the upper end of the hydrophobic pocket, or with Trp171, Arg92 and Tyr117 which are lower in the hydrophobic pocket and thereby place the CHL closer to the exit gate (see also Sup. Fig, 11A,B). An even lower position of the bound CHL was observed in the simulations (Fig. 1G), but it is visited only rarely because it triggers incipient water penetration from the aqueous environment which leads to energetically unfavorable interactions with the hydrophobic tail of CHL. Together, the three modes of CHL binding outline an energetically favorable path for the sterol in the functional steps of uptake and release when StarD4 is embedded in the membrane environment appropriate for these functions.

Indeed, our recent findings from a combined experimental/computational study (34) showed that StarD4 can preferentially extract sterol from liposome membranes containing certain PIPs (especially, PI(4,5)P_2_ and to a lesser degree PI(3,5)P2), while enhancement of transport was less effective when the same PIPs were present in the acceptor membranes. Moreover, we showed that StarD4 recognizes membrane-specific PIPs through specific interaction with the geometry of the PIP headgroup as well as the surrounding membrane environment.

Our key mechanistic finding of the connection between the movement of CHL among its binding modes in the hydrophobic pocket, and the movements of specific structural motifs of StarD4, emerged from the analysis of the very long MD trajectories with the time-resolved rare event detection algorithm RED (31). These trajectory analyses showed that the connected conformational changes at the β9 C-terminal and H4 N-terminal portions result in the opening of a gate between H4 and Ω1-loop, and a corridor between β9 and β7β8-loop. From the time-resolved RED analysis we were able to ascertain that these gate-opening/gate-closing conformational changes occur simultaneously with the transitions between CHL binding modes, with the opening corresponding to the position of CHL in the binding pocket near Trp171, Arg92 and Tyr117.

The opening of β8-β9 corridor and the partial unraveling of Helix4 N-terminal head (res199-201) in the Trp-binding state make the lower end of the hydrophobic pocket accessible to the external environment. Although being observed here directly for the first time, these function-related conformational changes are consistent with results from a previous study using a membrane penetration assay to study the environment of M206 (18). M206 is a residue located at the lower half of the hydrophobic pocket adjacent to the gate. In the crystal structure (PDBid:1JSS), it is facing inwards to the hydrophobic environment. However, in the absence of membrane, the iodoacetamide-NBD (IANBD) probe at the M206C site gave a signal similar to the L124C site immersed in water, which suggests that the M206 at the inner side of H4 is accessible to the external water environment. With the introduction of liposomes, the probe at the M206C site was found to be inserted into the nonpolar environment of the membrane. The demonstrated accessibility of M206 to the external water or membrane environments supports the inference that the Trp-binding state with opened gates and local unraveling in H4 represent an intermediate state of StarD4 in an aqueous environment that is also likely to be an intermediate state along the CHL release/uptake pathways when StarD4 becomes embedded in a target membrane.

Conformational changes of the H4-Ω1gate related to sterol release has long been proposed for StarD-protein family members (18,25,27,38), but the allosteric path and exact conformational changes remained unknown. While the Ω1 loop was suggested to be essential for the opening of the gate, the structural comparison analysis shown in (27,38) suggests as well a change in H4 – as seen in our study – likely as a result of steric interaction with the CHL at the Trp-binding site.

To understand the dynamics of the mechanism observed in the RED analysis, we employed the information-theory-based analysis (NbIT) (32,33) to quantify the allosteric interactions that connect the changes in CHL binding modes to the function-related conformational changes. The results identified the paths of allosteric connectivity and showed that the gating element H4 is indeed a strong allosteric coordinator of the cholesterol binding mode dynamics. The coordination is achieved dynamically as cholesterol moves to a binding site lower in the pocket, by (1) the straightening of the H4 helix that makes space for the movement of the CHL, and (2) H4 dissociating from the interaction with the Ω1 loop which opens the gate further. These coordinated changes trigger yet more conformational rearrangement as shown in macrostates 4 and 6 (Sup. Fig. 6B, 8D). Notably, this coordinated set of CHL translocation in the binding pocket and rearrangements of structural motifs result in a conformation similar to the structures of cholesterol-binding LAM protein complexes. There, the bound sterol is located close to the tunnel entrance rather than residing at the bottom of the hydrophobic cavity, and as the tunnel entrance opens slightly at the Ω1-loop, the ligand sterol is exposed to water (28–30). This similarity is illustrated in the structural superposition of StarD4 and LAM2 (Sup. Fig. 11C,D) where the location of the LAM ligand close to the entrance is shown to correspond to that of the CHL in the Trp-binding mode of StarD4, adjacent to the water network between W171, R92 and Y117 (39).

Notably, the allosteric network revealed by the NbIT analysis in the StarD4-cholesterol complex shows that the allosteric coordination between the cholesterol binding and the peripheral motifs includes the β1-3 and β9 sheets. The β1 and β2 sheets are known to be essential for StarD4 membrane-binding that is mediated by their basic residues patch which interacts with anionic lipids (18,34). Thus, we have shown recently (34) that upon membrane-embedding of StarD4 the β1 and β2 sheets bind PIP2 in a subtype-specific manner, which is likely to contribute to the recognition of target organelles based on the prevalent PIP2 subtype composition of their membranes. Also, membrane-embedded StarD4 adopts an orientation where the β9 and β7β8-loop motifs that are part of the allosteric network (Fig. 6), are interacting with lipid heads in the membrane (34).

Together, the results from the RED and NbIT analyses of the extensive set of long MD trajectories of cholesterol-bound complexes of WT StarD4 and the three mutants have revealed a well-defined allosteric mechanism in CHL-bound StarD4. The conformational rearrangements determined with the RED algorithm to occur simultaneously with the relocation of the CHL from the Ser-binding mode to the Trp-binding mode in the hydrophobic pocket are proposed to constitute a preparation for CHL release into a membrane environment. The mechanism of this preparation is defined in specific detail by the NbIT-quantified allosteric network of interactions between the CHL binding site and the peripheral structural motifs involved in cholesterol translocation. As β1 was reported to be an essential membrane-binding motif (18,34), we propose that when the bound StarD4 interacts with the membrane, the movement of CHL between binding modes propagate trough the allosteric pathway to change the conformations of the peripheral motifs of StarD4 to enable delivery. We are currently evaluating this hypothesis in membrane-embedded StarD4 systems described in (34).

## MATERIALS AND METHODS

### Modeling of the cholesterol StarD4 complex

The structure of mouse apo-StarD4 was taken from the X-ray crystal structure (PDB ID: 1jss), which includes residues 24 to 222 of the protein (17). Lys223 and Ala224 were introduced using Modeller 9.23 software (40), resulting in the conformation of mouse StarD4 22-224 segment with N-terminus acetylation. StarD4 complexes with cholesterol (CHL) were obtained by docking the ligand in the hydrophobic pocket of StarD4 using Schrodinger’s Induced Fit Docking protocol (35,36). The common features shared by the top 15 docking results are: the interaction of the cholesterol hydroxyl head that forms 2 or 3 hydrogen bonds with the sidechains of Ser136 & Ser147 (Fig. 1A) and with the backbone carbonyl of Cys148, and the sequestering of the cholesterol hydrocarbon tail inside the hydrophobic pocket, away from the water environment. This top docking mode was used as the initial structure for the MD simulations of the StarD4-CHL complex solvated in 0.15M K^+^Cl^-^ ionic aqueous solution (∼32,600 atoms).

### Long unbiased simulation of cholesterol-bound StarD4 in water

The cholesterol-bound StarD4 system was equilibrated first in MD simulations with NAMD version 2.12 (41) using a multi-stage protocol. In the first stage the backbone of StarD4 and the heavy atoms of the ligand were harmonically constrained and gradually released in three steps of 0.5ns each, with restraining force constants changing from 1 kcal/(mol·Å2) to 0.5, and 0.1, respectively. This stage was followed by 6ns unbiased MD simulation using NAMD. In the third (production) stage, the velocities of all the atoms were reset, and the system was simulated with openMM software (42) in 12 independent replicates, each for 33.6μs, resulting in a cumulative simulation time of 403μs (>0.4 milliseconds). The OpenMM simulations were conducted in NPT ensemble (T=310K, P=1 atm) using a 4fs integration time-step and a Monte Carlo Membrane Barostat.

### Mutant constructs of the StarD4-CHL complex

Mutations were introduced into the StarD4-CHL complex with the free-energy perturbation protocol in NAMD version 2.12 (41). Accordingly, hybrid StarD4 models containing overlapped residues (e.g., overlapped K49 and A49 to introduce the K49A mutation) were constructed using representative conformations obtained in the most popular Ser-binding state 1 and Trp-binding state 3. Then, the mutations are introduced by gradual annihilation of the interactions between the WT residue and its surrounding, achieved by decreasing the coupling parameters from 1 to 0 in 10 steps, and concomitantly increasing the interactions between the mutant residue and its surrounding from 0 to 1 in 10 steps. The simulation was run for 5ns per window. After the last window, the annihilated original residues are removed from the hybrid StarD4 models, and the mutant StarD4 is equilibrated for another 10ns. For production runs, 24 replicas are made for each system and sampled in long unbiased simulation with openMM (42).

### Rare Event Detection (RED) Protocol

Rare events in MD trajectory data were detected as described with our RED protocol (31) which utilizes a Non-negative Matrix Factorization (NMF) algorithm implemented in the Scikit-Learn python package (43).The unsupervised machine learning algorithm learns to decompose a trajectory into a set of components that represent structural motifs that move simultaneously, and reports their appearance as dominant structural characteristics during a particular period in the course of the trajectory. Briefly summarized, the event detection method in the RED protocol uses the NMF algorithm to learn a sparse, parts-based representation of the data (44). For trajectory analyses the input array (I) is a contact matrix constructed as described below from the data in each frame of the trajectory. By construction, this array encodes the information about the protein structure as it evolves over the trajectory time. The NMF algorithm decomposes the I array into a “spatial” array and a “temporal” array, which together are responsible for the integrated detection of the temporally defined rare events that provide the mechanistic information. Thus, the columns of the “spatial” output matrix produced by the NMF analysis are termed “components”, as each of them represents a set of conformational features that change simultaneously even if they are in different structural motifs. As the determinant features of a conformational state evolve over trajectory time, the different components containing them dominate the structural characteristics of the molecule at different times. Thus, the corresponding temporal array represents the contribution of each component along the trajectory, and thus identifies the time period where a particular component dominates the structural characteristics of the protein (31). A simple illustration provided in the full description of the method (31) refers to the analysis of a conformational transition in the unfolding of a fully folded alpha helical segment. The analysis results in one component that is the folded structure, and a second one that is the unfolded structure. Each frame of the simulation trajectory will be a mixing of these archetypes, with the folded component dominating the trajectory frames at the time segments before the transition, and the unfolded component dominating those after the transition.

#### Building the time-evolution contact map for the input array (**I**)

A contact map is constructed between every pair of residues for each frame in the trajectory, with contact marked as 1 if any atom of one residue is < 3.5Å away from any atom in another residue. For the 201 residues of StarD4 this yields a tensor of 0 and 1 values with dimensions (t, 201,201), where t represents the number of frames. Subsequently, the tensor is rearranged to a matrix of size 201^2^ columns and t rows, where 201^2^ is the total number of residue pairs. The matrix columns, representing all the residue pairs, are further trimmed by excluding the contact pairs that remain invariant throughout the entire simulation, which yields a ***I*** matrix with n_pairs=3880 columns. Each column represents a contact pair that has altered its contact state at least once during the entire simulation, encapsulating protein dynamics information for further analysis.

To focus on the conformational changes around the time of cholesterol relocation between its binding modes in the hydrophobic pocket, we excerpted 6 trajectory segments from the 6 independent runs as indicated by the time intervals (**traj0**, from 6.5 to14.5μs; **traj1**, 0.1-4.1μs**; traj3**, 0.1-8.1μs; **traj4**, 0.1-6.5μs; **traj6**, 0.1-4.1μs; **traj7**, 0.1-4.1μs; 34.4μs total). In order to obtain results that are generalizable among trajectories (which decreases the sensitivity to incidental structural fluctuations and benefits the reproducibility of RED analysis), the contact matrices of the 6 segments are inputted at the same time as the training data for the NMF algorithm. This input of multiple trajectories is constructed by concatenating the data matrices along the time-axis. This contact map is recorded every 0.8ns, which yields an ***I*** matrix with t=43000 rows. Smoothing is applied to the contact matrix across the time dimension by averaging the array in a sliding window of 20 nanoseconds (25 frames).

#### Details of the protocol

The algorithm decomposes the input contact matrix described above into multiple components, where each component has a structural state described by a “spatial” row array with length n_pairs (here n_pairs=3880), and a corresponding “temporal” column array of length t (here t=43000). The number of components (c) is a super-parameter chosen by the user according to the number of independent processes in the system and was set to be c=5 to study generalizable features shared in independent runs. Thus, the ensemble of the “temporal” column arrays is a temporal matrix of size (t, c) in this study, and the ensemble of the “spatial” row arrays is a spatial matrix of size (c, n_pairs=3880), where the multiplication of the temporal matrix and the spatial matrix is fitted to resemble the time-evolution contact map data matrix. Each row in the c=5 rows of components in the spatial matrix is an array that encodes the determinant conformational features that evolve concurrently over time. The corresponding column in the temporal matrix represents the contribution of these features in determining the conformational state of the protein at each time. It and identifies the trajectory times at which this particular set of features dominates the structural characteristics of the protein by showing it to have the highest “weight” in this period of trajectory time.

#### Interpretation of the results

As shown in Fig. 2E-G, the spatial arrays scatter plots show that there are groups of pairs that form a horizonal line, suggesting that these pairs are constantly contributing similarly to every component. These groups of pairs that are salient in every component by being in contact throughout the StarD4 simulation, and contribute similarly to each component, are used for normalization of the spatial arrays. Thus, each spatial array is divided by a normalization factor that equals to the mean value of the constructive residue pairs contributing to this particular component, thereby placing the contact features of these constitutive residue pairs at 1 in Fig. 2E-G and Sup. Fig. 2B. To preserve the relative weight between components, each corresponding temporal array is multiplied by the same normalization factor derived from the spatial array of this particular component, so that the multiplication result of spatial and temporal matrix remains the same and resembles the input contact matrix.

This normalization allows for direct comparison between components. Subtracting the spatial array of comp A from comp B, as represented in (Sup. Fig. 3), cancels out the contributions of the constitutive residue pairs so that pairs with positive contributions are those newly established in the event corresponding to the AèB transition, whereas the negatively contributing pairs are those breaking contact during this AèB event (Sup. Fig. 3). These pairs are salient in the transition as they contribute to the structural differences between components and are termed *structure differentiating contact pairs* (SDCPs). The SDCPs that are more salient than 4 times the standard deviation, are summarized in Table 1 and are grouped based on the motif movement they represent.

### Dimensionality reduction with the “time-structure based independent component analysis” (tICA) approach

The MD trajectory frames were projected onto a space of 7 collective variables described in Sup. Table 1, with the “time-structure based independent component analysis” (tICA) method. The tICA method identifies the slowest reaction coordinates of a system in the dynamic dataset (45–47) by solving for the generalized eigenvectors and eigenvalues of the time-lagged covariance matrix:

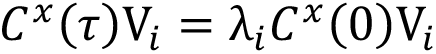

in which the *C^x^*(τ) is the time-lagged covariance matrix *C^x^*(τ) = 〈𝑋(𝑡)𝑋^*T*^(𝑡 + 𝜏)〉_$_ between collective variables, 𝐶^*x*^(0) = 〈𝑋(𝑡)𝑋^*T*^(𝑡)〉_$_ is the regular covariance matrix, 𝑋(𝑡) is the data vector at time 𝑡, 𝜏 is the lag-time, *V_i_* is the i^th^ tICA eigenvector, and 𝜆_*i*_ is the i^th^ tICA eigenvalue. In this study, the stride of the trajectories is 0.4 ns/frame, and the lag-time 𝜏 for the covariance matrix construction is 10ns (25 frames), thus ignoring any fluctuations faster than 10ns which are not relevant to the slow processes in the system. The first 1μs in every independent run are discarded to equilibrate the systems.

### Construction of the Markov State Model (MSM) and assignment of Macrostates

#### 1. Construction of the Markov State Model

Markov State Model (MSM) is a powerful tool to study the kinetics of equilibrium state-transition processes in protein function or activation pathways (48–50). The implementation and validation of MSMs are well documented (51–53). To construct MSMs, the 3D conformational space of the first three tIC vectors was discretized into microstates. The transition count matrixes between the microstates were then recorded in a transition count matrix, which is then symmetrized by averaging with its transpose matrix in order to satisfy detailed balance and local equilibrium (53,54). Finally, the transition probability matrixes (TPMs) were built by normalizing the transition count matrix by the transitions departing from the same microstate.

#### 2. Selection of parameters for Markov model construction

The choice of super parameters: the lag-time of the transition, and the discretization of the conformational space (i.e., number of microstates), are optimized to ensure the quality of MSM (53,54). The lag-time is optimized with implied timescale test:

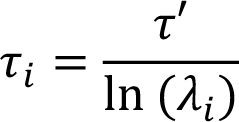

in which 𝜏′ is the lag-time used for building the transition matrix, 𝜆_*i*_ is the i^th^ eigenvector of the TPM, and 𝜏_*i*_is the implied timescale corresponding to the i^th^ relaxation mode of the system. In this study, the lag-time of 120 ns defined as shown in Sup. Fig. 5A ensures the best available Markovian behavior with the minimal loss of data. The best discretization of the conformational space is chosen by comparing the resulted MSM with the generalized matrix Rayleigh quotient (GMRQ) method (54,55), where the MSM is trained on a randomly picked training set comprised of half the set of trajectories, and then cross-validated on the test set which includes the other half of the trajectory set. The method tests how well the eigenvectors of the training set diagonalize the TPM of the test set, scored as:

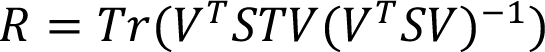

in which 𝑇𝑟 denotes trace of the inner matrix product, 𝑉 contains the first n eigenvectors of the training set, 𝑆 is the diagonal matrix composed of equilibrium populations of the test set (i.e. a diagonal matrix composed by the first eigenvector of the test set TPM), and 𝑇 is the test set TPM. The best MSM parameters yield the highest GMRQ score. In this study, the 200 k-means microstates were chosen so as to provide a good balance between the consistency shown by GMRQ result (Sup. Fig. 5B) and the coverage of the 3D conformational space.

#### 3. Interpretation of MSM, and Macrostate assignment

To extract information from the MSM, TPM is decomposed into its eigenvalues and eigenvectors. The largest eigenvalue is 1, and its corresponding first eigenvector represents the equilibrium state population of the system. The remaining eigenvectors represent the population flow between microstates. The 2^nd^ largest eigenvalue corresponds to the eigenvector representing the population flow contributing to the slowest relaxation process of the system. In turn, the next largest eigenvalues and eigenvector correspond to the next slowest relaxation process, and so on. Here, the 200 microstates were grouped into 9 macrostates according to their kinetics similarities learnt from MSM, using the using the Robust Perron Cluster Analysis (PCCA+) algorithm (56).

### Transition path theory (TPT) analysis

The transition paths among the 9 macrostates were obtained from the TPT analysis (57), which is based on the flux matrix between states whose components are defined as:

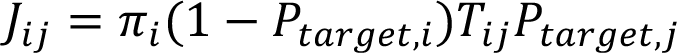

where 𝜋_*i*_ is the equilibrium population of state I, 𝑇_*ij*_ is the transition probability from state i to state j, and 𝑃_*target,i*_ is the probability of visiting the target state i as the first state during the transition before visiting other states. The most probable transition pathways between specific states was obtained by iteratively finding the next pathway with the highest flux in the flux matrix using a graph theory algorithm, the Dijkstra algorithm (58), implemented in the MSM builder software package (54).

### Quantifying allosteric effect using N-body Information Theory (NbIT)

#### **1.** NbIT analysis focused on multiple motifs in the StarD4-CHL complex

The information theory-based NbIT analysis was used to measure the coordination between distanced motifs and to quantify the allosteric effect (32,33). For the calculation of the coordination information we chose the following structural motifs: **CHLsite:** S136 S147 W171 R92 Y117; **β1**: res R46 V47 A48 K49 K50 V51 K52; **β2β3**: res R58 K59 P60 Y67 L68 Y69; **β2β3loop**: E62 E63 F64 N65; **H4head**: Q199 S200 A201 D203 T204 A207; **Ω1loop**: L124 N125 I126; β9: D192 R194 G195; **β7β8loop-nearβ9**: V162 R163 G164; **β7β8loop-mid**: T157 R158 P159 E160; **β7β8loop-nearβ6**: E153 W154 S155 E156; **H4tail**: R218 K219 G220 L221; **β3tail**: G73 V74 M75 D76; **β8β9loop**: S179 P180 S181 Q182. The allosteric effect was then quantified from the coordination information as described in ref 32.

Briefly, the first step in the calculation of the coordination information was the alignment of the trajectories to a reference structure – we used the StarD4 crystal structure (PDB ID: 1JSS). The alignment of snapshot (per 0.4ns) was carried out on the alpha carbons of residues in β3-9 and H4, excluding the motifs have been found to undergo conformational changes (70-76, 79-86, 103-108, 113-118, 131-140, 146-150, 170-175, 184-189, 211-220). With the aligned trajectory, the entropy in each motif was calculated analytically through the differential entropy:

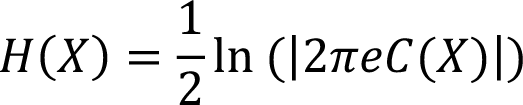

in which 𝐶(𝑋) is the covariance matrix describing all variables in X, and X is the displacement array describing the position of every atom in the motif by its distance to the ref conformation. Then, the *Total Correlation* (TC) describing the total amount of information share in a set of motifs was calculated as:

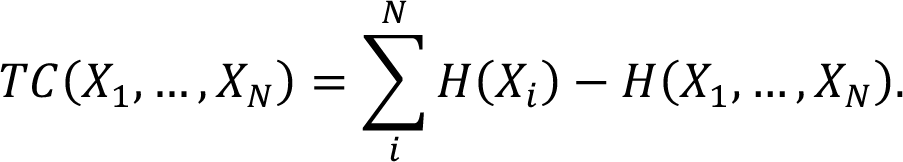

And the *Coordination Information* (CI) describing the amount of information shared in the receptors that is also shared with another motif that works as the *transmitter*:

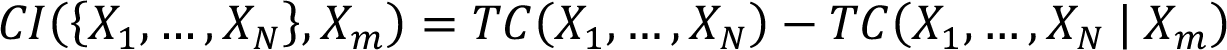

Here 𝑇𝐶(𝑋_1_, …, 𝑋_-_|𝑋.) is the conditional total correlation between {𝑋_1_, …, 𝑋_-_} conditioning on 𝑋*_m_*:

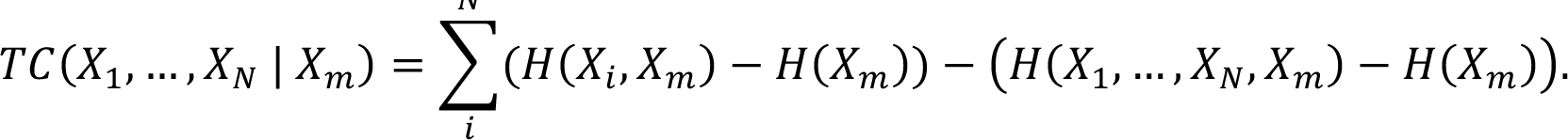

The *Coordination Information* was then normalized by the *Total Correlation* of the *receiver*, to represent the portion of dynamics of the receptor that is allosterically coupled with the *transmitter*.

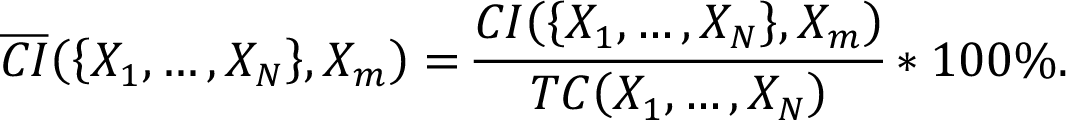

In Table 3 and the Supplementary Tables 2, 4, 5, and 6, the *Total Correlation* (TC) within each motif is displayed along the diagonal, while the *Normalized Coordination Information* is presented in the off-diagonal elements, with residues on the top (columns) acting as the *Transmitter* and residues on the left (rows) being the coordinated *Receiver*.

#### 3. Coordination channel analysis based on mutual coordination information

To define the allosteric channels that mediate the coordination information (Fig. 6A), we calculated the *mutual coordination information* that measures the amount of coordination information shared between the *receiver* and the *transmitter* motifs <u>that is also shared</u> with another structural element:

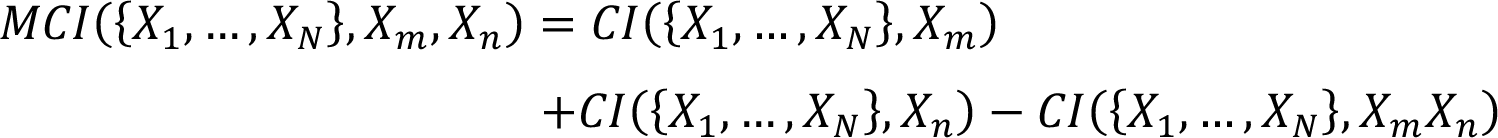

Here 𝑋.𝑋_/_ is the union set constituted by the residues in the *Transmitter* 𝑋. and the *channel* 𝑋_/_. The *mutual coordination information* was then normalized by the *Coordination Information* between the receiver and the *transmitter* to obtain the NMCI values used in the determination of the allosteric pathway in Fig. 6B.

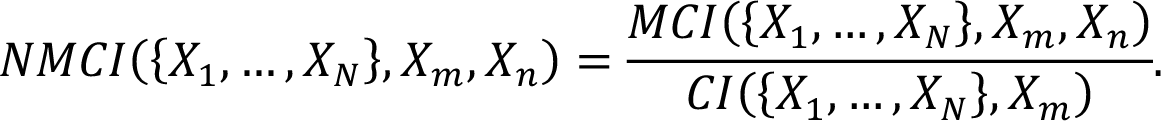

## ACKNOWLEDGEMENTS

We gratefully acknowledge helpful discussions with Drs. Ambrose Plante, Ekaterina D. Kots, and Prof. George Khelashvili. HX gratefully acknowledges their helpful guidance regarding the principles, applicability, implementation, and interpretation of the RED, NbIT and TPT analysis. We are grateful for discussions with Prof. Frederick R. Maxfield and his lab members. Support from the 1923 Foundation for the project “*How Needed Molecular Precision is Achieved for Addressing, Pickup, and Delivery in the Trafficking of Cholesterol Among Cell Membranes*” is gratefully acknowledged. The computational resources and technical help at the Center for Computational Innovations (CCI) at the Rensselaer Polytechnic Institute (RPI), and the efficient and sustained access to the AiMOS supercomputer at CCI generously awarded through the COVID-19 High Performance Computing Consortium, are gratefully acknowledged. We are grateful for the computational resources under Projects BIP225 and BIP109 at the Oak Ridge Leadership Computing Facility, which is a DOE Office of Science User Facility supported under Contract DE-AC05-00OR22725, and for the in-house computational resources of the David A. Cofrin Center for Biomedical Information in the Institute for Computational Biomedicine at Weill Cornell Medical College.

## Data availability

Atomistic MD simulations were carried out with OpenMM 7.5. Computational analysis was carried out using a combination of python scripts, and in-house scripts available on GitHub https://github.com/weinsteinlab/. Computational data presented in the manuscript will be managed in full accordance with the institution’s Data Management and Sharing policy of Weill Cornell Medical College which is in full compliance with the NIH requirements. The raw data used to reach the inferences and conclusions are available upon reasonable request from HW.

