## Supplementary Figures and Tables for "Allosterically coupled conformational dynamics in solution prepare the sterol transfer protein StarD4 to release its cargo upon interaction with target membranes"

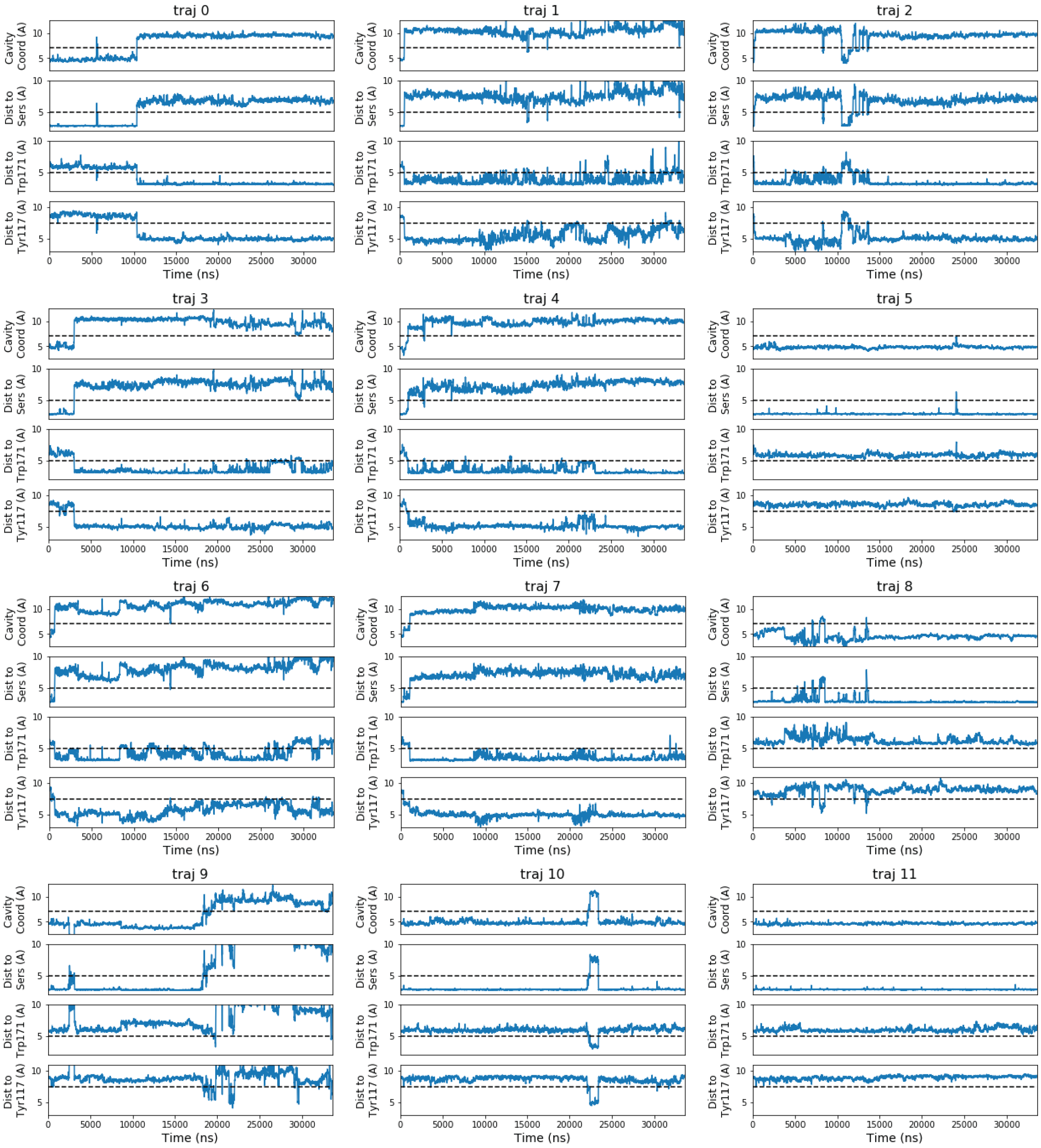


**Supplementary Figure 1: Time-evolution of cholesterol binding modes in each trajectory.** Each trajectory is represented by a **stack of four panels** showing the time evolution of a binding mode quantification. Increases in Y axis values indicate progress towards the bottom of the cavity, close to the gate**. Upper panel:** the cholesterol “cavity coordinate” (see definition of the cavity coordinate measure in Fig. 1B). **Second panel**: the distance between the cholesterol oxygen to the sidechain oxygen of the nearest Ser-binding site (S136, S147). **Third panel:** the distance between the cholesterol oxygen to the nitrogen in W171 sidechain. **Fourth panel:** the distance from the cholesterol oxygen to the sidechain oxygen of Y117.

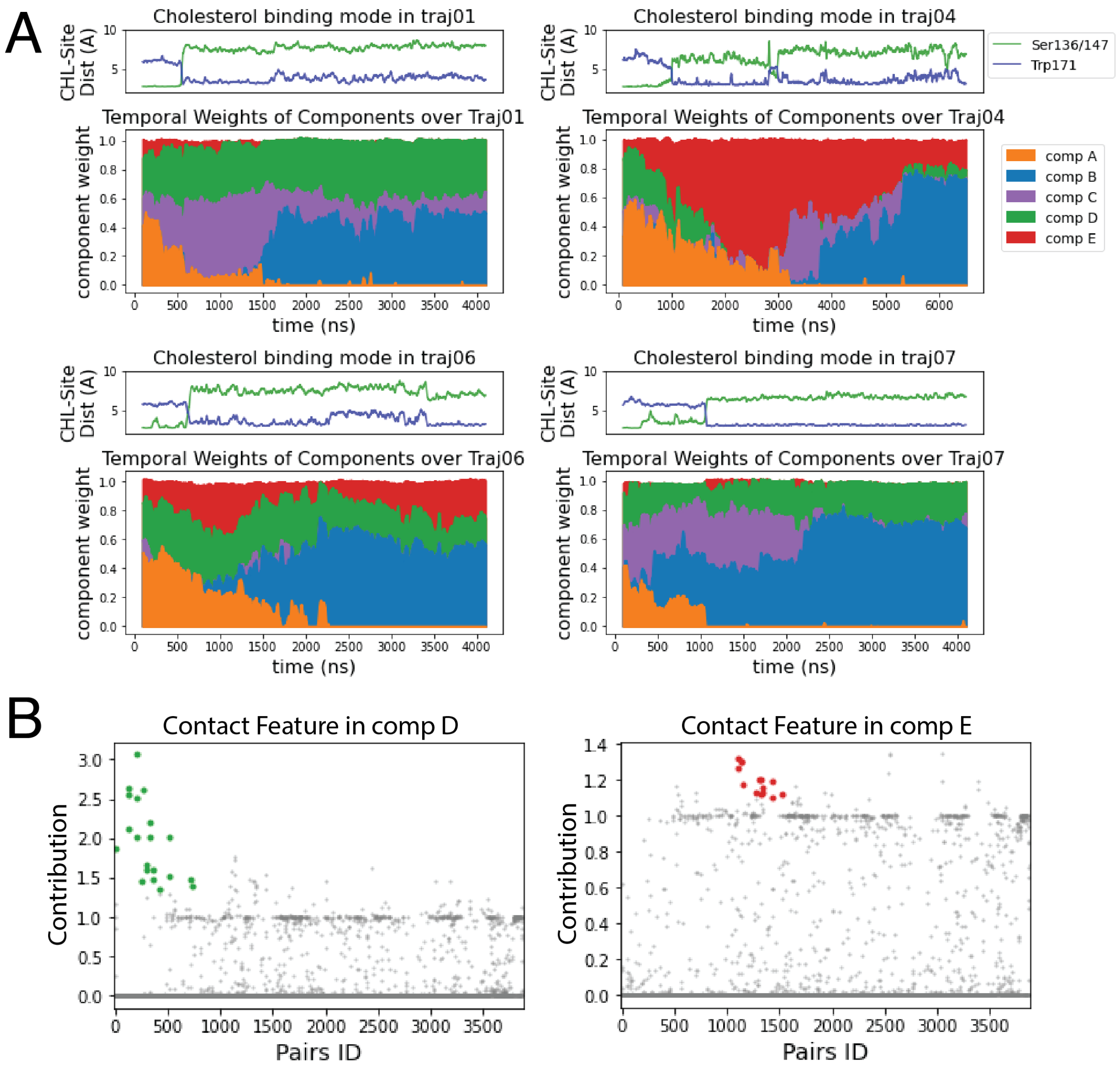


**Supplementary Figure 2: Detailed analysis of RED results: temporal weights and spatial arrays** (compare to Fig. 2 in main manuscript). **(A)** Relation between time evolution of the cholesterol binding mode (Upper panel) and the RED-detected temporal weight of structural components (solid color panels) over the simulation time of trajectories 1 (top left panel); 4 (top right); 6 (bottom left); and 7 (bottom right). In the Upper panel the evolution of the cholesterol binding mode is represented by the distance from CHL to Ser site (in green) and to Trp site (in blue) plotted over the simulation time. In the Temporal Weights panels, the normalized temporal weights of the RED components are identified by color over the simulation time: comp A (orange), comp B (blue), comp C (purple), comp D (green), comp E (red). At each time point, each position represents *the stacked* value of all components beneath, as explained in Fig. 2A (e.g., the weight of comp A in traj01 is near 0 at 2000, while comp B is ~0.44, comp C is ~0.20, and comp D is ~0.37). **(B)** Normalized spatial arrays of components D (left) and E (right) show the contact features between residue pairs. Along the X axis there are 3880 data points each representing a residue pair, and the Y coordinate of each data point shows the contact feature of the residues pair captured by the component (see the Methods). Structural differentiating residue pairs are highlighted in colors and presented in Sup. Fig. 3C-F.

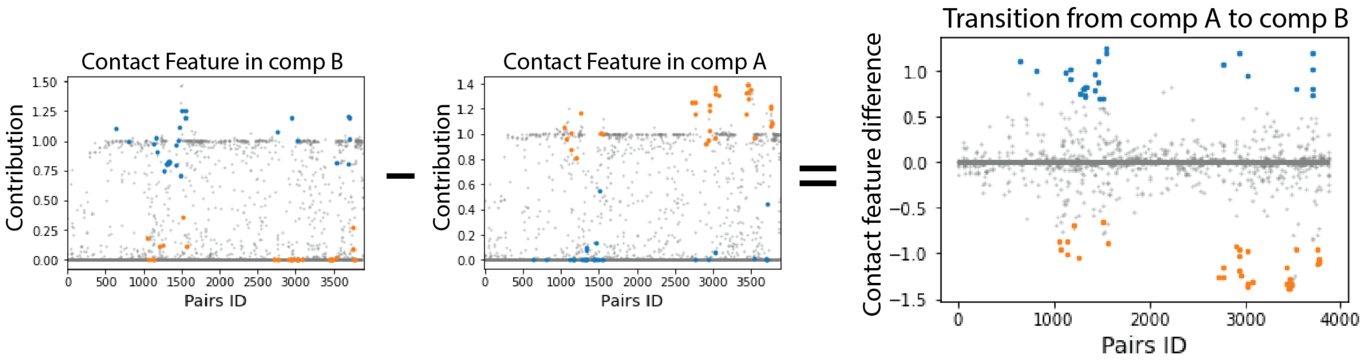


**Supplementary Figure 3: Comparison of normalized spatial components reveals the contact breaking and forming during a conformational state transition.** A contact feature difference diagram (far right) obtained by subtracting the corresponding elements of the spatial array of comp A from those of the comp B array identifies the pairs contributing to the transition event. The positively contributing pairs are those with a high contact rate in comp B but a low contact rate in comp A, i.e, pairs that **gain** contact in the transition from comp A to comp B (blue dots in the difference diagram). The pairs shown by orange dots are the negatively contributing pairs that were in contact only in comp A and **lost** contact when transitioning from comp A to comp B.

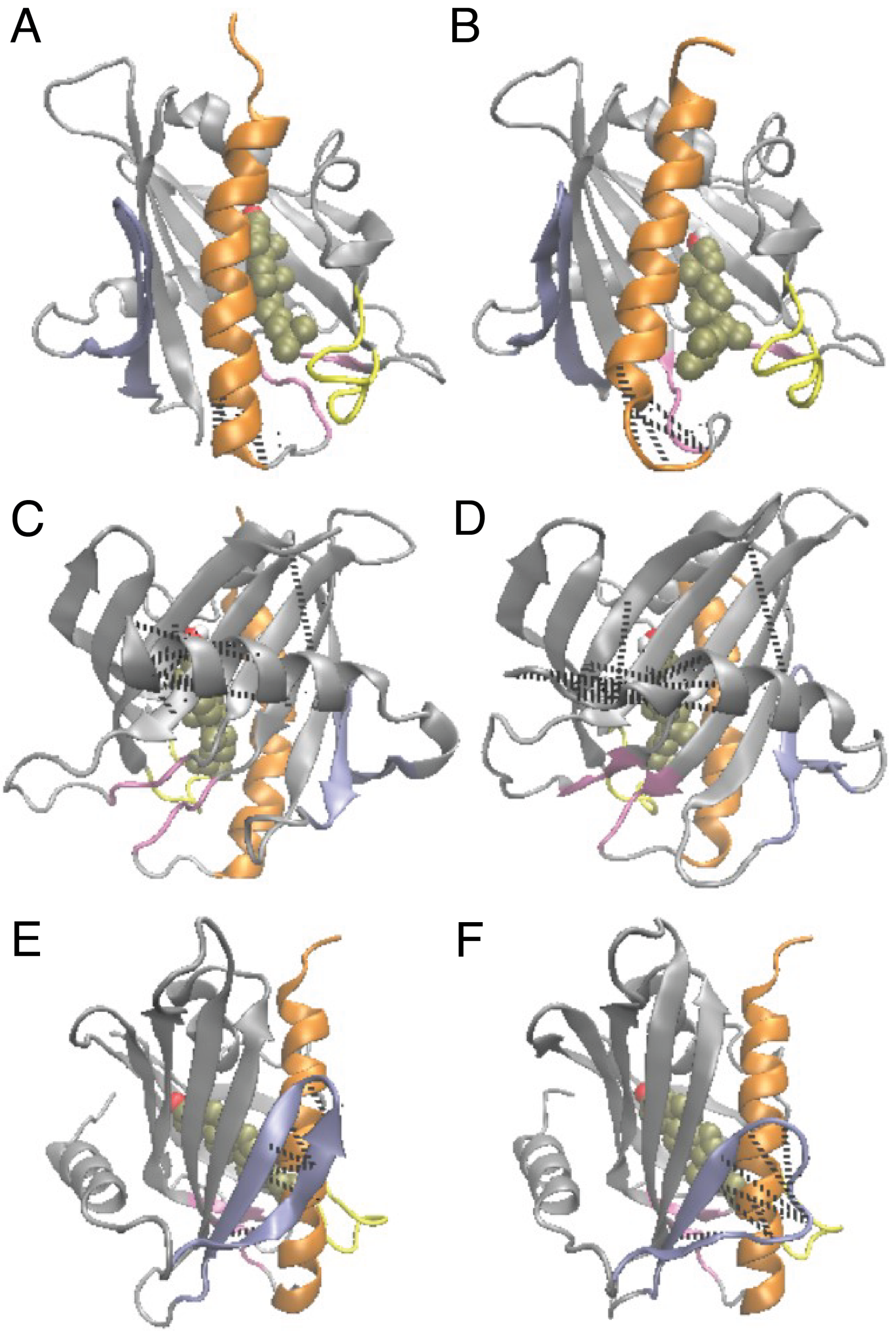


**Supplementary Figure 4. Structural elements reported by RED to be involved together in conformational changes but are not coupled with cholesterol movement. (A,B)** Residue pairs in group 5 are labeled with dashed lines in panel **A** showing the N-terminal head on the bottom of Helix4 in the folded state **(A)** and in the state shown in panel **B,** which is a partially unfolded state. **(C,D)** Structures that emerged in Component D: Partial unfolding of the Helix1 N-terminal head is shown in the folded state **(C)** and the unfolded state **(D)**. The residue pairs that are salient in comp D (Sup. Fig. 2B) are labeled with dashed lines. **(E,F)** Structures that emerged in Component E: The unfolding of the β1 is sheet shown in the folded state **(E)** and the unfolded state **(F)**. The salient residue pairs in comp E (Sup. Fig. 2B) are labeled with dashed lines.

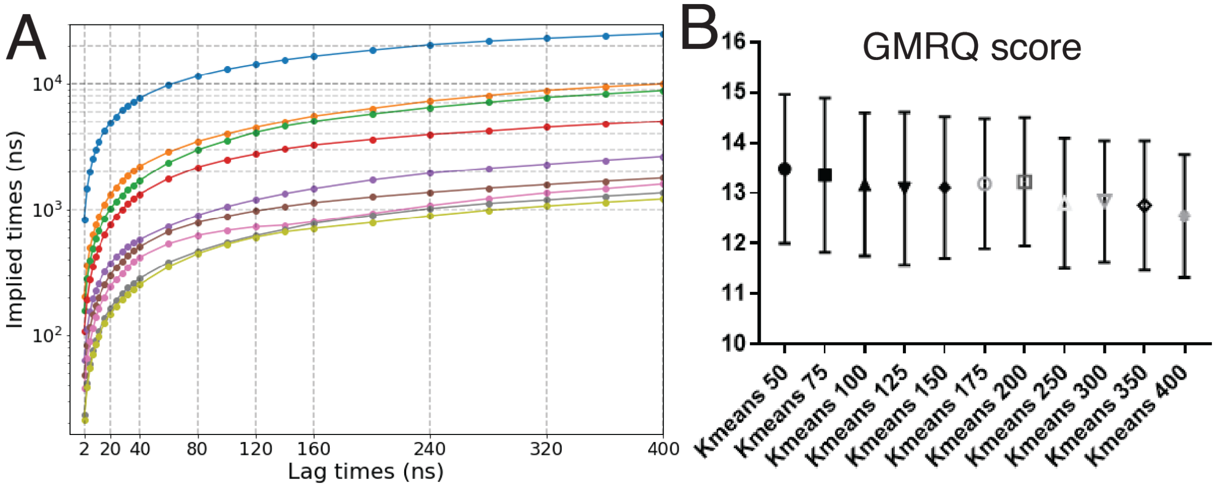


**Supplementary Figure 5. Quality checks in the construction of the Markov State Model.**
**(A)** Implied timescales calculated for the 9 slowest relaxation modes in the system. The implied timescales reach a plateau after lagtime >300ns, with no change in the relative sequence of the 9 slowest relaxation modes. **(B)** GMRQ scores examining the conformational space discretization methods: conformational space is discretized into 50 to 400 states using k-means clustering. Error bars are standard deviations calculated from 500 GMRQ tests, obtained from randomly generated training sets and test sets, each containing half of the trajectories.

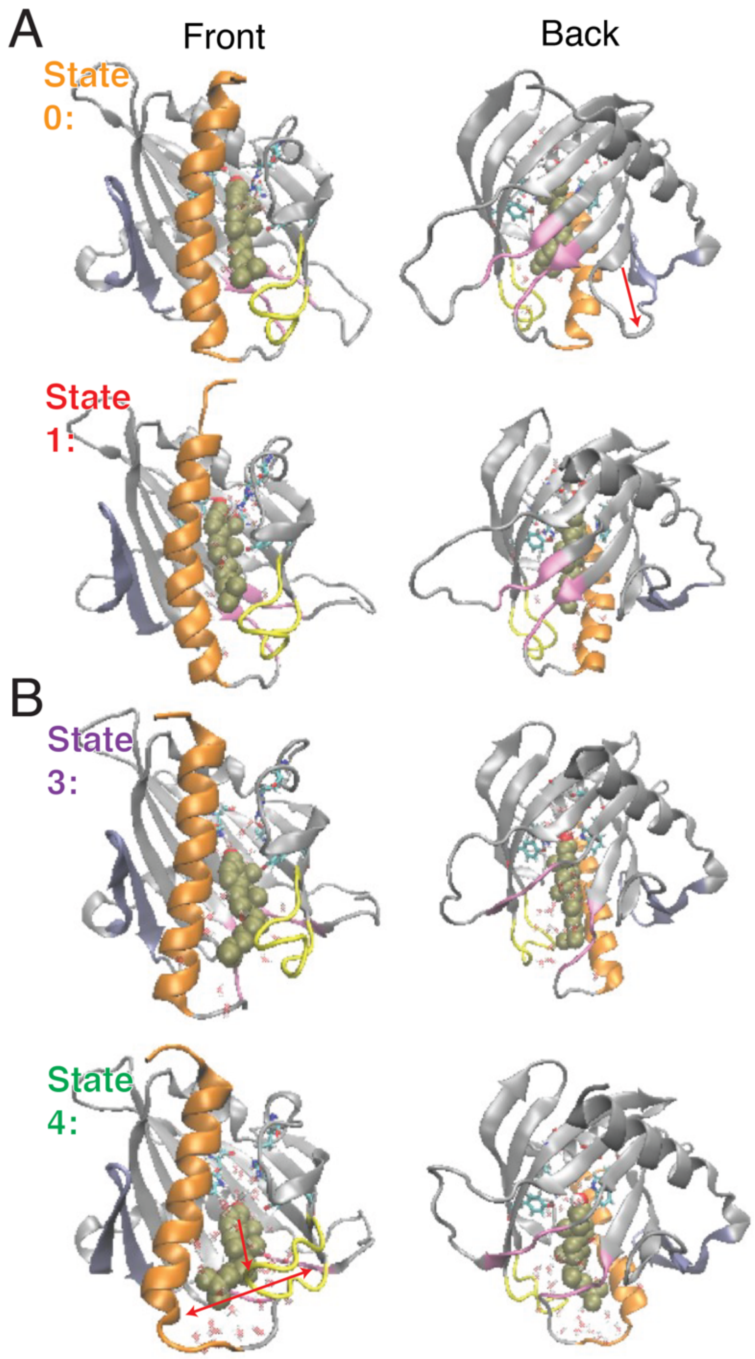


**Supplementary Figure 6. Representative conformations for macrostates 0 and 4, shown in comparison to representative conformations for macrostates 1 and 3.** Representative models for macrostates 0, 1 **(in A)** and macrostates 3, 4 **(in B)** are viewed from front and back as labeled. StarD4 is rendered in gray, with the C-terminal Helix (Helix4) in orange, the loop between β5 and β6 (Ω1 loop) in yellow, the β1&2 sheets in blue, and the β8 loop and β9 loop in pink. CHL is rendered in VDW with hydroxyl group in red and white and other atoms in tan color. Residues S136, S147, W171, R92, Y117 are rendered in "licorice", as are the water molecules within 6Å from CHL. Red arrow in the 1st row labels the conformational change at β23loop of the structure of state 0 in comparison to state 1. Red arrows in the State 4 structure label the conformational change at H4-Ω1 gate of the structure in comparison to state 1.

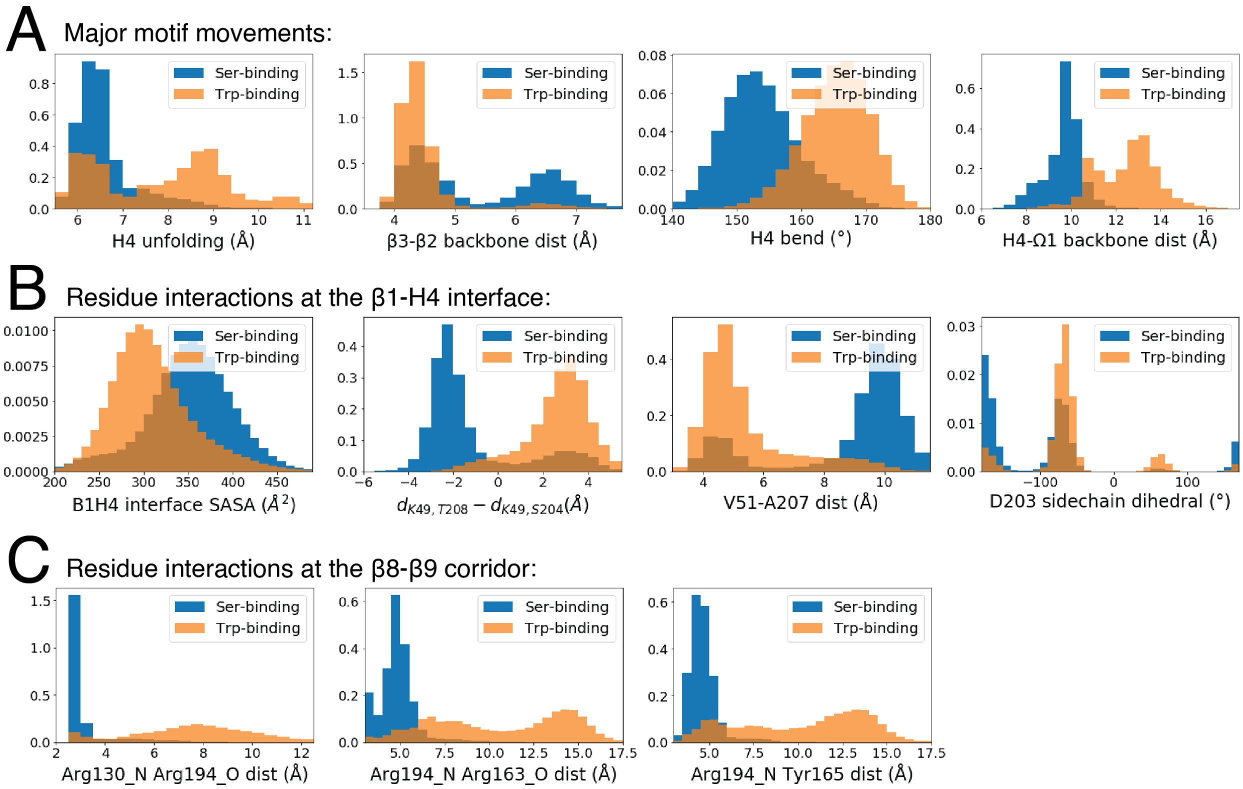


**Supplementary Figure 7: (A-C) Histograms of CVs showing the changes occurring in StarD4 structural elements during the conformational transition event detected by RED.** Comparison of probability distributions of the CV values in the representative structures of macrostate 0 and 1 in which CHL is in the Ser-binding mode (blue histograms), and in macrostate 3 where CHL is in the Trp-binding mode (orange histograms). **(A)** CVs capturing major motif movements. **(B)** CVs capturing the residue interactions at the β1-H4 interface. **(C)** CVs capturing the residue interactions at the β8-β9 corridor. The CVs are defined in Sup. Table 3.

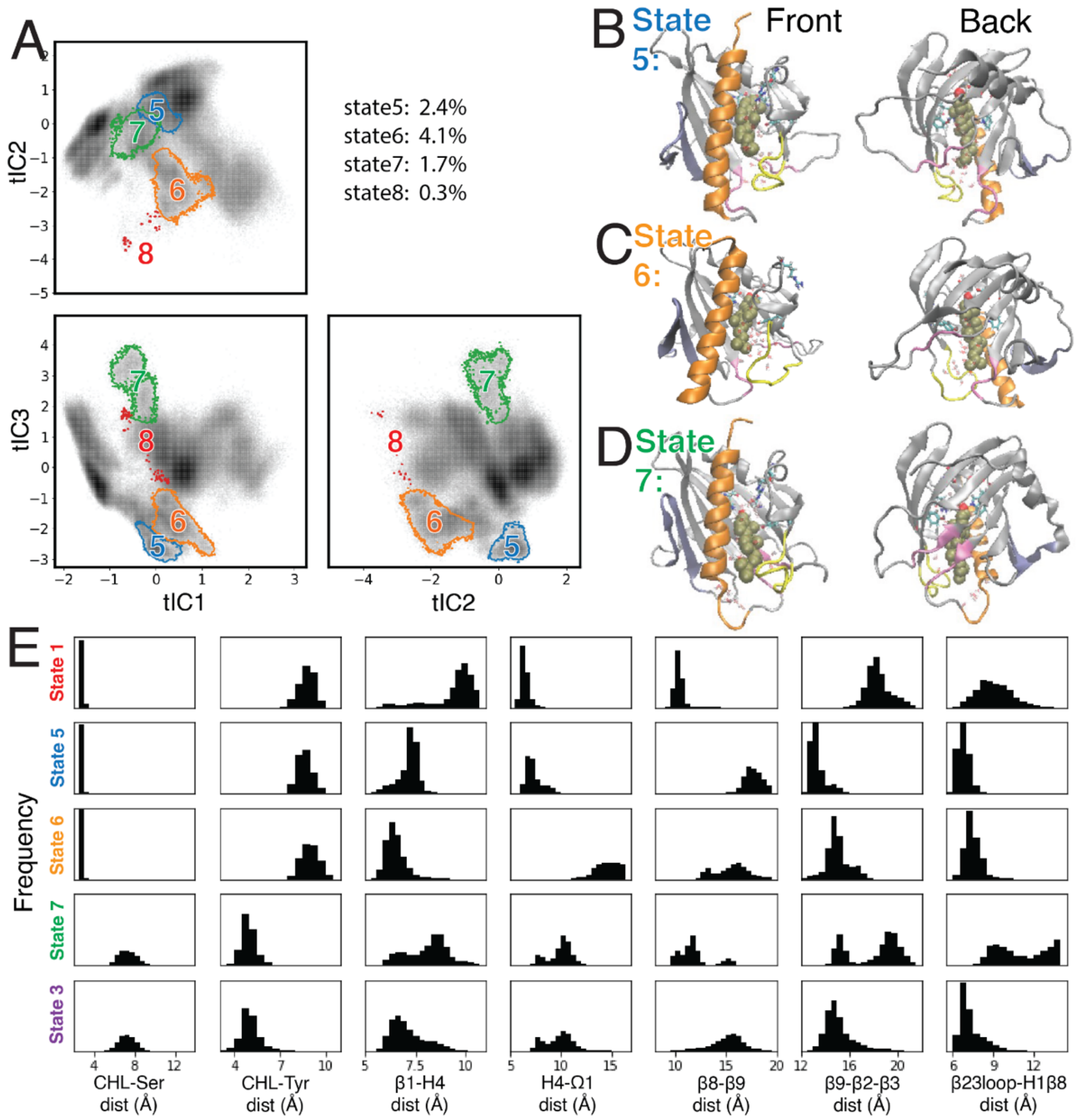


**Supplementary Figure 8: Same analysis as in Figures S6 and S7 for representative structures of the lower populated (intermediate) macrostates in which CHL translocation and gate opening are not coupled. (A)** Macrostates 5 (blue), 6 (orange), 7 (green), 8 (red) are outlined in the 3D conformational space, projected on the tIC2-tIC1 plane (upper left), tIC3-tIC1 plane (lower left), tIC3-tIC2 plane (lower right). The gray shade represents the population density. The populations in these macrostates are listed at the upper right. **(B)** Structural model representing macrostate 5 in which CHL is in the Ser-binding mode with the H4-Ω1 gate closed but the β8-β9 corridor open. **(C)** Structural model representing macrostate 6 in which CHL is in the Ser-binding mode but both the H4-Ω1 gate and the β8-β9 corridor open widely. **(D)** Structural model representing macrostate 7 in which CHL is in the Trp-binding mode with the H4-Ω1 gate open but the β8-β9 corridor closed. StarD4 is rendered in gray, with Helix4 in orange, the loop Ω1 loop in yellow, and the β1β2sheets in blue. CHL is rendered in VDW, and water molecules within 6Å from CHL are shown in transparent licorice. **(E)** Structural characteristics of the macrostates 1, 3, and 5 to 7 6 shown as the probability density histogram of the characteristic CVs. For comparison, the histograms are grouped (1,5,6); (7,3) by similarity. The CVs are defined in Sup.Table1.

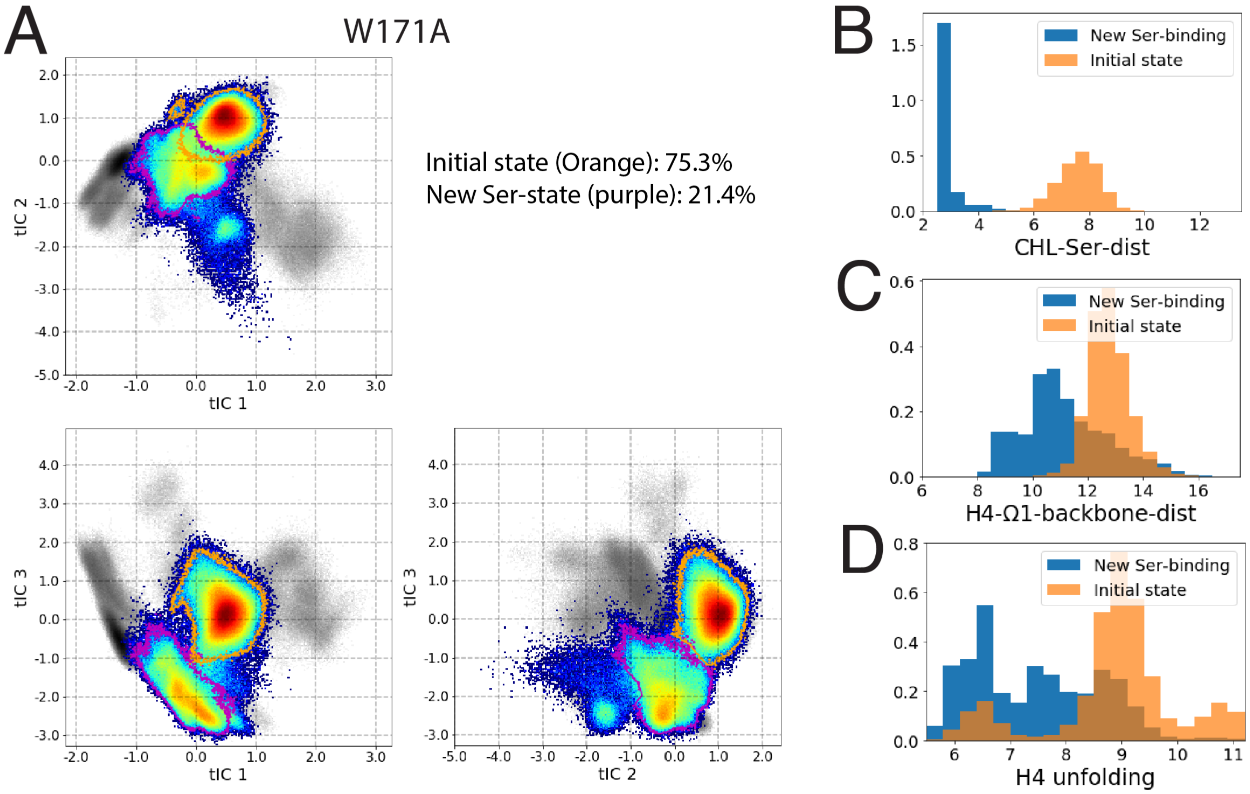


**Supplementary Figure 9: Conformation analysis of W171A StarD4 dynamics. (A)** The 3D conformational space covered by the trajectories of W171A StarD4 simulations started from the Trp-binding-like state. The gray shade represents the population density of the original conformational space sampled by WT StarD4, while the colored map represents the population density of the W171A StarD4 construct as in Fig. 4C. Only two major states were sampled in the W171A StarD4 simulations, and the populations are listed at the upper right. **(B-D)** Histograms of CVs showing the changes occurring in structural elements of W171A StarD4 during the simulations of conformational transition between the two major states. Results for the starting Trp-binding-like state are in orange; results for the Ser-binding state resulting from the transition of CHL to the other binding mode, are in blue. The CVs are **(B)** CHL-Ser-dist, defined in Sup. Table 1; **(C)** H4-Ω1-backbone-dist, defined in Sup. Table 1; and **(D)** H4 unfolding, defined in Sup. Table 3.

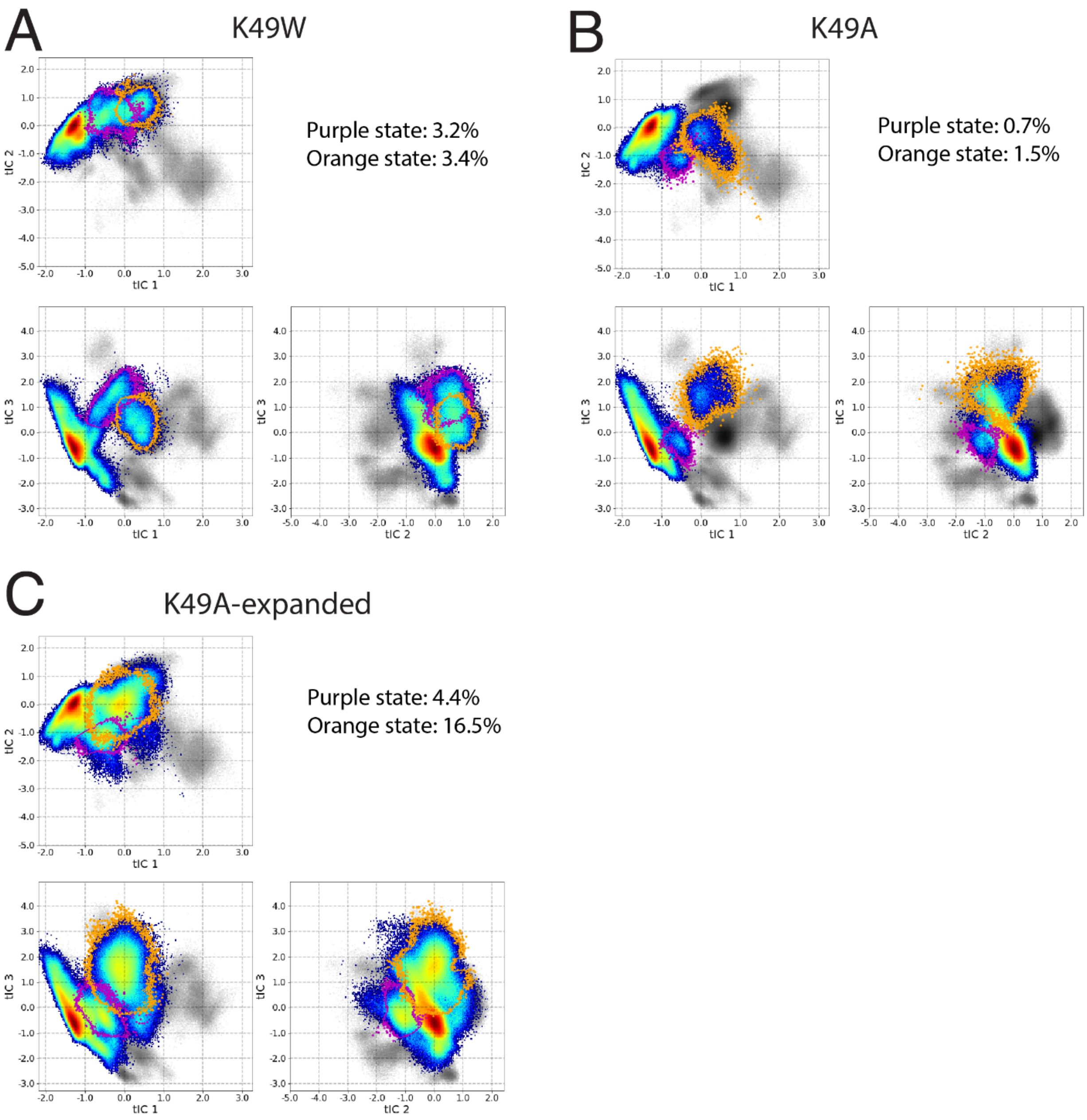


**Supplementary Figure 10: The 3D conformational tICA space covered by the trajectories of K49 mutants of StarD4 starting from the Ser-binding-like state: (A)** Simulations of K49W StarD4; **(B)** The first batch of K49A StarD4 simulations; **(C)** The expanded K49A StarD4 simulations. The gray shade represents the population density of the original conformational space sampled by WT StarD4, while the colored maps represent the population density of the mutant StarD4 systems (see Fig. 4C for details). Two new metastable states were sampled in each conformational tICA space, and the populations are listed at the upper right.

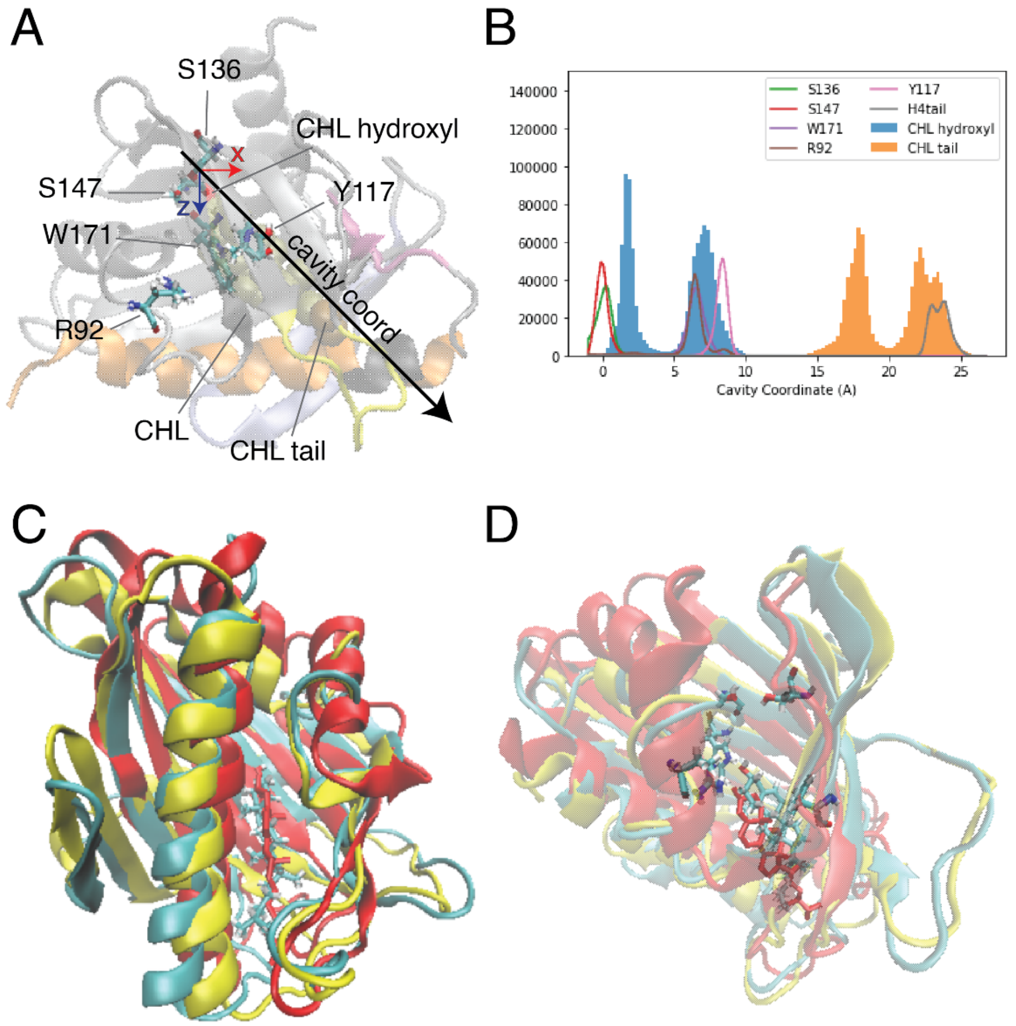


**Supplementary Figure 11. Binding position of the CHL ligand relative to the residues in the hydrophobic pocket and compared to the position of the CHL ligand in the crystal structure of LAM2** (PDB ID 5YS0)**. (A)** The motifs of are interest labeled in a conformation of StarD4 rendered in the same way as in Fig. 1A, in an orientation where the "H4 tail" residues 202 to 205 are in the back. The CHL binding site residues S136, S147, R92, W171, Y117 are rendered in "licorice". The CHL is rendered in VDW with the oxygen in red and the carbon atoms C25, C26, C27 at the tail colored in ochre. The "cavity coordinate" is defined in Fig 1B. **(B)** Frequency histograms of the cavity coordinate for: the CHL oxygen (filled blue), CHL tail (filled orange). The hollow line plots show the distributions of the following structural elements identified by the color code insert: the sidechain oxygen or nitrogen atoms in S136, S147, W171, R92 and Y117, and of the center of mass of the H4 tail. **(C,D)** Structural superposition of StarD4 on the crystal structure of LAM2. In red: the crystal structure of the second StARkin domain of Lam2 in complex with ergosterol (PDB ID 5YS0), with the sterol ligand shown in red licorice. In *yellow*: the crystal structure of mouse StarD4 with no ligand (PDB ID 1JSS). In *cyan*: a representative conformation of CHL-StarD4 complex at the Trp-binding state, with the sterol ligand and the binding site residues shown in licorice.

**Supplementary Table 1. The definition of collective variables (CVs) used as tICA parameters**

| **CVs** | **Definitions** |
|---|---|
| CHL-Ser-dist (Å) | The minimal distance from the cholesterol oxygen to the sidechain oxygens in the Ser136/147 binding site |
| CHL-Tyr-dist (Å) | The distance from the CHL oxygen to the sidechain oxygen of Tyr117 * |
| β1-H4-dist (Å) | The mean distance between the backbone of residue pairs on β1 and H4: K49-T204, V51-A207 |
| H4-Ω1-pairwise-dist (Å) | The mean distance between residue pairs that form hydrophobic interactions between H4 and Ω1: Q122-A205, L123-A201, I127-A201, L123-A205, I127-A205 |
| β6-β9-dist (Å) | The distance between the Center of Mass (COM) of the backbone of residues 192 and 195 on β9, and the COM of the backbone of residues 130 to 133 on β6 ** |
| β9-β3-β2-dist (Å) | The sum of distances: the distance between the COM of the backbone of residues 58 to 61 on β2 and COM of the backbone of residues 190 to 194 on β9, and distance between the COM of the backbone of the residues 58 to 61 on β2 and the backbone of the residues 66 to 68 on β3 |
| β23loop-H1β8-dist (Å) | The mean distances between residue pairs from β2β3 loop to H1 or β8: Y37-F64, F64-H167, F64-P168 |

* The distance of CHL to the Trp-binding site is measured using Tyr117 as the reference position instead of Trp171, since CHL may move pass Trp171 while Tyr117 is the lowest site as shown in Fig 1C

** The distance of β9 to β8 is measured using β6 as the reference position instead of β8, since β9-β6 distance presents clearer movement of β9 than using the more flexible β7β8loop as the reference)

**Supplementary Table 2. Normalized coordination information in non-correlated motifs serve as negative controls**

Coordinator:

Receiver:

| | H4tail | β3tail | β8β9 loop | CHLsite |
|---|---|---|---|---|
| **H4tail** | 5.79 | 5.8% | 3.0% | 6.0% |
| **β3tail** | 11.5% | 3.51 | 13.6% | 10.3% |
| **β8β9loop** | 2.6% | 9.8% | 6.68 | 3.6% |
| **CHLsite** | 14.2% | 15.8% | 11.5% | 1.70 |

**Supplementary Table 3. The definition of CVs used in detailed structural analysis**

| CVs | Definitions |
|---|---|
| **H4 unfolding (Å)** | The sum of distances between the backbone of residue pairs on the first turn of H4: Q199-D203, S200-T204, and A201-A205 |
| **H4 bend (°)** | The angle between two vectors: the vector from the Cα of residue 208 to the Cα of residue 202, and the vector from the Cα of residue 210 to the Cα of residue 216 |
| **β3-β2-dist (Å)** | The distance between the CoM of the backbone of residues 65 to 68 on β3 and the CoM of the backbone of residues 58 to 61 on β2 |
| **H4-Ω1 backbone dist (Å)** | The distance between the CoM of the backbone of residues 122 to 127 on Ω1 and the CoM of the backbone of residues 201 to 205 on H4 |
| **β1H4 interface SASA (Å2)** | The solvent accessible surface area contributed by residues A48 K49 V51 on β1 and residues D203 T204 A207 S208 A211 on H4 |
| **$d_{K49,S208} - d_{K49,T204}$ (Å)** | Distance difference: the distance from the CoM of the sidechain of K49 to the CoM of the sidechain of S208, **minus** the distance from the CoM of the sidechain of K49 to the CoM of the sidechain of T204 |
| **V51_A207_dist (Å)** | The distance from the CoM of the sidechain of V51 to the CoM of the sidechain of A207 |
| **D203 sidechain dihedral (°)** | The dihedral angle of the atom Cγ, Cβ, Cα and N of D203 |
| **Arg130_N Arg193_O dist (Å)** | The distance from the sidechain nitrogen of R130 to the backbone oxygen of R194 |
| **Arg194_N Arg163_O dist (Å)** | The distance from the sidechain nitrogen of R194 to the backbone oxygen of R163 |
| **Arg194_N Tyr165 dist (Å)** | The distance from the sidechain nitrogen of R194 to the sidechain aromatic ring of Y165 |

**Supplementary Table 4. Normalized coordination information between sites in cholesterol-StarD4 W171A system**

Transmitter:

| Receiver: | β1 | β2β3 | β2β3 loop | H4 head | Ω1 loop | β9 | β7β8loop-nearβ9 | β7β8loop-mid | β7β8loop-nearβ6 | CHLsite |
|---|---|---|---|---|---|---|---|---|---|---|
| **β1** | 15.78 | 21.9% | 9.6% | 15.3% | 6.9% | 6.8% | 6.5% | 7.7% | 6.1% | 8.4% |
| **β2β3** | 29.4% | 10.71 | 29.5% | 21.1% | 11.6% | 11.7% | 9.1% | 10.9% | 9.2% | 13.9% |
| **β2β3loop** | 15.6% | 34.8% | 7.22 | 12.9% | 8.4% | 8.7% | 8.2% | 11.2% | 9.7% | 11.8% |
| **H4head** | 16.2% | 21.2% | 10.5% | 11.97 | 10.8% | 13.4% | 8.4% | 9.2% | 7.8% | 9.6% |
| **Ω1loop** | 8.0% | 8.7% | 6.4% | 11.5% | 6.30 | 8.1% | 9.0% | 9.6% | 7.2% | 8.6% |
| **β9** | 8.1% | 12.3% | 9.7% | 21.4% | 8.8% | 3.32 | 7.9% | 7.7% | 6.3% | 6.5% |
| **β7β8loop-nearβ9** | 7.1% | 6.6% | 5.5% | 7.3% | 9.5% | 7.1% | 3.67 | 33.6% | 11.2% | 7.3% |
| **β7β8loop-mid** | 6.9% | 7.4% | 7.0% | 7.8% | 7.0% | 6.4% | 20.1% | 9.81 | 34.0% | 8.1% |
| **β7β8loop-nearβ6** | 6.4% | 9.6% | 8.9% | 9.2% | 6.7% | 9.8% | 16.4% | 33.5% | 7.40 | 8.3% |
| **CHLsite** | 40.1% | 41.3% | 37.2% | 39.6% | 28.0% | 27.9% | 31.7% | 36.9% | 32.8% | 1.55 |

* The normalized coordination information (NCI) is presented with residues on the top (columns) acting as the *Transmitter* and residues on the left (rows) being the coordinated *Receiver*. On the diagonal, the total correlation (TC) of the site is shown in gray.

**Supplementary Table 5. Normalized coordination information between sites in cholesterol-StarD4 K49W system**

Transmitter:

| Receiver: | β1 | β2β3 | β2β3 loop | H4 head | Ω1 loop | β9 | β7β8loop-nearβ9 | β7β8loop-mid | β7β8loop-nearβ6 | CHLsite |
|---|---|---|---|---|---|---|---|---|---|---|
| **β1** | 10.95 | 16.3% | 8.4% | 13.1% | 6.1% | 6.0% | 4.6% | 7.4% | 7.7% | 9.2% |
| **β2β3** | 19.2% | 8.22 | 26.4% | 14.3% | 7.9% | 8.4% | 6.3% | 8.5% | 8.1% | 10.9% |
| **β2β3loop** | 8.6% | 29.3% | 7.84 | 8.0% | 7.0% | 5.4% | 4.6% | 6.9% | 6.5% | 9.7% |
| **H4head** | 13.9% | 17.4% | 8.3% | 10.60 | 7.9% | 11.2% | 5.5% | 8.6% | 9.1% | 10.4% |
| **Ω1loop** | 6.1% | 7.7% | 6.9% | 9.0% | 5.34 | 7.2% | 6.2% | 6.5% | 5.7% | 8.0% |
| **β9** | 5.5% | 9.1% | 7.2% | 16.4% | 6.2% | 3.47 | 6.3% | 5.4% | 4.6% | 7.2% |
| **β7β8loop-nearβ9** | 3.8% | 6.5% | 4.8% | 5.0% | 8.4% | 9.6% | 2.27 | 15.9% | 6.0% | 7.9% |
| **β7β8loop-mid** | 6.6% | 7.1% | 6.2% | 6.5% | 4.8% | 5.5% | 12.6% | 10.23 | 39.3% | 7.9% |
| **β7β8loop-nearβ6** | 6.3% | 6.6% | 5.7% | 6.5% | 4.1% | 5.0% | 12.9% | 35.1% | 7.71 | 7.9% |
| **CHLsite** | 35.6% | 38.5% | 33.1% | 35.7% | 30.0% | 24.2% | 27.7% | 34.1% | 34.8% | 1.44 |

* The normalized coordination information (NCI) is presented with residues on the top (columns) acting as the *Transmitter* and residues on the left (rows) being the coordinated *Receiver*. On the diagonal, the total correlation (TC) of the site is shown in gray.

**Supplementary Table 6. Normalized coordination information between sites in cholesterol-StarD4 K49A system**

Transmitter:

| Receiver: | β1 | β2β3 | β2β3 loop | H4 head | Ω1 loop | β9 | β7β8loop-nearβ9 | β7β8loop-mid | β7β8loop-nearβ6 | CHLsite |
|---|---|---|---|---|---|---|---|---|---|---|
| **β1** | 16.19 | 29.6% | 11.1% | 23.1% | 4.7% | 13.9% | 7.9% | 8.5% | 6.9% | 9.8% |
| **β2β3** | 30.2% | 12.17 | 30.3% | 18.1% | 4.2% | 15.3% | 8.6% | 9.2% | 7.1% | 10.8% |
| **β2β3loop** | 8.1% | 26.9% | 11.28 | 9.1% | 1.5% | 9.6% | 5.8% | 6.2% | 4.5% | 5.4% |
| **H4head** | 21.1% | 22.0% | 12.0% | 17.21 | 5.1% | 20.3% | 10.7% | 11.3% | 7.8% | 11.1% |
| **Ω1loop** | 2.7% | 3.3% | 2.2% | 5.3% | 6.63 | 5.8% | 1.9% | 2.3% | 1.9% | 2.9% |
| **β9** | 16.1% | 21.5% | 16.4% | 25.1% | 5.8% | 6.78 | 14.7% | 12.7% | 10.2% | 13.8% |
| **β7β8loop-nearβ9** | 4.6% | 8.6% | 6.4% | 10.0% | 2.1% | 7.2% | 7.42 | 35.1% | 12.9% | 8.2% |
| **β7β8loop-mid** | 6.2% | 8.6% | 6.1% | 9.3% | 2.3% | 8.2% | 23.7% | 12.70 | 33.8% | 7.9% |
| **β7β8loop-nearβ6** | 5.7% | 6.9% | 5.4% | 7.5% | 2.7% | 6.8% | 13.6% | 37.7% | 9.32 | 7.2% |
| **CHLsite** | 18.8% | 23.3% | 15.4% | 19.3% | 10.1% | 18.7% | 24.6% | 22.3% | 19.7% | 1.58 |

* The normalized coordination information (NCI) is presented with residues on the top (columns) acting as the *Transmitter* and residues on the left (rows) being the coordinated *Receiver*. On the diagonal, the total correlation (TC) of the site is shown in gray.
